# Relationships of Worldwide *Onobrychis* species: from seed & pod morphology and molecular perspective

**DOI:** 10.1101/2022.05.02.490301

**Authors:** Mohammad Zarrabian, Azam Nikbakht Dehkordi, Mohammad Hossein Ehtemam, Mohammad Mahdi Majidi

## Abstract

The present study aimed to investigate the phylogeny of 31 *Onobrychis* species with the help of seed and fruit macro & micro-morphological traits and ISSR molecular markers. The species were found to be highly variable in their seed and fruit morphological traits. Also, a new type of fruit (*O. crista* galli type), in addition to the three predefined types of fruit, was introduced. From molecular perspective, a set of 22 ISSR primers were shown to have a good efficiency for genetic discrimination among the species. Bayesian model based STRUCTURE analysis grouped the genotypes into 4 genetic clusters, and the high value of Fst calculated for these clusters indicated a high level of genetic differentiation among them. On the other hand, based on the Maximum Likelihood phylogenetic tree, our results are in agreement with the subgenus classification (*Onobrychis* and *Sisyrosema*), while contrary to *Sisyrosema*, the *Onobrychis* subgenus (the *Lophobrychis* and *Onobrychis* sections) appears not to be monophyletic. Furthermore, in contrast to the traditional taxonomic classification, species belonging to the *Lophobrychis* section do not form a coherent group, indicating that this section could be considered as a heterogeneous unit and perhaps could be merge into the *Onobrychis* section.

**Highlight:** This paper discusses a worldwide phylogenic analysis and population structures of *Onobrychis* genus. The results of this paper provide a comprehensive answer to complexity of subgenus and section discrimination.

## Introduction

The genus *Onobrychis Mill.,* which belongs to the Fabaceae family, includes about 170 species around the world. This genus is widely distributed in northern temperate regions of the world whereas Iran and Turkey are considered to be the center of diversity for this species. (Rechinger, 1969; Yildiz *et al*., 1999; Hayot-Carbonero, 2011). This genus is divided into two subgenera: *Onobrychis* and *Sisyrosema,* which are distinguished from each other through deciduous flowers, glabrous corollas, erect fruits, and the epidermis of a calyx with crystals within (Yildiz *et al*., 1999). Moreover, subgenus *Onobrychis* includes four sections (*Dendrobrychis* DC, *Lophobrychis*, *Onobrychis*, and *Laxiflorae*) while the subgenus *Sisyrosema* comprises five sections (*Anthyllium*, *Afghanicae*, *Heliobrychis*, *Hymenobrychis*, and *Insignes*) (Rechinger, 1969).

Up until now, several approaches have been adopted to provide an appropriate classification of the *Onobrychis* genus. Yildiz *et al*. (1999) recognized 170 species based on fruit morphological characters and re-evaluated the sectional delimitation of the *Onobrychis* genus. However, later studies suggested that it would be necessary to re-examine the taxonomic position of the *Onobrychis* taxa (Cenci *et al*., 2000; Abou-el-Enain, 2002). A karyological study within *Onobrychis* clade found that sub-genus *Sisyrosema*, unlike subgenus *Onobrychis*, is monophyletic (Hejazi *et al*., 2010). Arslan and Ertugrul (2010) maintained that a greater similarity could be established on the basis of seed storage protein between the *Heliobrychis* and *Hymenobrychis* sections rather than the *Onobrychis* section.

Unlike studies on protein and characteristics of seeds, DNA fingerprinting revealed that a classification of the *Onobrychis* genus weakly supported the taxonomical sections and many species would be synonyms (Hayot-Carbonero, 2011; Safaei Chaei Kar *et al*., 2014; Duan *et al*., 2015; Zarrabian and Majidi, 2015; Amirahmadi *et al*., 2016). In the previous nrDNA ITS-based phylogenetic studies (Ahangarian *et al*., 2007; Safaei Chaei Kar *et al*., 2012), the non-monophyly of the *Onobrychis* subgenus indicated the subgenus *Sisyrosema* was retained as monophyletic. On the other hand, Amirahmadi *et al*. (2014) suggests that none of these two subgenera is monophyletic. More studies based on three non-coding chloroplast sequences and the nuclear ribosomal DNA internal transcribed spacer revealed that section *Heliobrychis* was retrieved as a well-supported monophyletic group sister to *Hymenobrychis* (Kaveh *et al*. 2018). It seems that the challenges and uncertainties are the results of the multitude of recognized species and the high variability of morphological characters due to their wide geographical dispersion (Zarrabian *et al*., 2013). To expand our knowledge on classification and genetic structure of the *Onobrychis* genus, the present study was conducted with the following objectives: 1) To illustrate the structures of the fruit (legume) surface and the external morphology of mature seeds, 2) To evaluate the diagnostic value of the investigated characters in terms of their systematic implications, and 3) To clarify phylogenetic relationships within the genera.

## Methods and martial

### Plant material

Eighty-eight accessions belonging to 31 species of the genus *Onobrychis* were used in this study (Table 1). The germplasms were obtained from the Leibniz Institute of Plant Genetics and Crop Plant Research (IPK), United States Department of Agriculture (USDA), and some species were collected from different geographical regions nationwide by the authors. All the accessions used for molecular experiments were germinated and grown at 25 °C with a 16/8 h day/night cycle in the greenhouse.

**Table 1.** Information of species and accessions investigated in this study

|  |  |  |  |  |  |
| --- | --- | --- | --- | --- | --- |
| Num. | Sub-genus | Section | Species | Number | Origin |
| 1 | *Onobrychis* | *Onobrychis* | *O. transcaucasia* | 9 | Armenia, Iran, Turkey, Georgia and Uzbekistan |
| 2 | *Onobrychis* | *Onobrychis* | *O. arenaria* | 10 | Soviet Union, Russia, Romania and Azerbaijani |
| 3 | *Onobrychis* | *Onobrychis* | *O. iberica* | 3 | Soviet Union and Pakistan |
| 4 | *Onobrychis* | *Onobrychis* | *O. cyri- Grossh* | 2 | Russia |
| 5 | *Onobrychis* | *Onobrychis* | *O. altissima* | 7 | Soviet Union, Russia, Azerbaijani, Georgia and Iran |
| 6 | *Onobrychis* | *Onobrychis* | *O. alba* | 3 | Bulgaria and Soviet Union |
| 7 | *Onobrychis* | *Onobrychis* | *O. inermis* | 4 | Russia |
| 8 | *Onobrychis* | *Onobrychis* | *O. petraea* | 5 | Germany, Iran and Russia |
| 9 | *Onobrychis* | *Onobrychis* | *O. oxyodonta* | 1 | Soviet Union |
| 10 | *Onobrychis* | *Onobrychis* | *O. gracilis* | 1 | Bulgaria |
| 11 | *Onobrychis* | *Onobrychis* | *O. persica* | 1 | Iran |
| 12 | *Onobrychis* | *Onobrychis* | *O. peduncularis* | 1 | Spain |
| 13 | *Onobrychis* | *Onobrychis* | *O. hajastana* | 1 | Soviet Union |
| 14 | *Onobrychis* | *Onobrychis* | *O. megataphrose* | 1 | Turkey |
| 15 | *Onobrychis* | *Onobrychis* | *O. montana* | 1 | France |
| 16 | *Onobrychis* | *Onobrychis* | *O. viciifolia* | 10 | Iran, China, USA, Czech Republic, Kyrgyz Republic, Spain, Ukraine, England, Morocco, Russia |
| 17 | *Onobrychis* | *Onobrychis* | *O. kemulariae* | 1 | Soviet Union |
| 18 | *Onobrychis* | *Lophobrychis* | *O. caput-galli* | 3 | Turkey, Israel and Unknown |
| 19 | *Onobrychis* | *Lophobrychis* | *O. crista-galli* | 4 | Iran, Israel and Unknown |
| 20 | *Onobrychis* | *Lophobrychis* | *O. aequidentata* | 3 | France |
| 21 | *Onobrychis* | *Lophobrychis* | *O. pulchella* | 1 | Turkmenistan |
| 22 | *Sisyrosema* | *Heliobrychis* | *O. melanotricha* | 2 | Iran |
| 23 | *Sisyrosema* | *Heliobrychis* | *O. argyrea* | 1 | Turkey |
| 24 | *Sisyrosema* | *Hymenobrychis* | *O. ptolemaica* | 2 | Iraq and unknown |
| 25 | *Sisyrosema* | *Hymenobrychis* | *O. hypargyrea* | 2 | Turkey |
| 26 | *Sisyrosema* | *Hymenobrychis* | *O. michauxii* | 2 | Turkey and Iran |
| 27 | *Sisyrosema* | *Hymenobrychis* | *O. chorassanica* | 1 | Soviet Union |
| 28 | *Sisyrosema* | *Hymenobrychis* | *O. sintenisii* | 3 | Soviet Union and Iran |
| 29 | *Sisyrosema* | *Hymenobrychis* | *O. bobrovii Grossh* | 1 | Russia |
| 30 | *Sisyrosema* | *Hymenobrychis* | *O. pallasii* | 1 | Soviet Union |
| 31 | *Sisyrosema* | *Hymenobrychis* | *O. radiata* | 1 | Armenia |

### Seed Morphological analysis

For seed Macro-morphological observations, seven seeds and legumes were placed under a binocular stereoscopic microscope (Osk 1844-b) (Table S1&S2). Moreover, fifteen seeds and legumes of each accession were measured for dimensions using a Digital Caliper (Osk 9888) and their average was used for each species (Table S1 and Fig. S1). Also, mature seeds of the species under investigation were selected for SEM study. For this purpose, the seeds were mounted on stubs using a double-face carbon tape and covered with gold (Au) at 15 mA for 3 min under high vacuum in an Ion Sputter Coating Unit (Baltek SCD-005). The samples were subsequently detected and photographed using SEM (Philips XL 130). The terminology of Botanical Latin Stearn (Stearn, 1983; Barthlott,1984; and Punt et al.,1994) was used to describe the SEM aspects of seed coat ornamentations (Figure S2&S3). For the cross-section of seed coats, three seeds were randomly selected from each accession, and the seed coats were separated before they were soaked in distilled water for 45 min. For convenience and uniformity of the cuts, a razor blade was used for making cross-sections. Transverse sections were cleaned with sodium hypochlorite for 1 min and slides were examined under a Nikon ECLIPSE E600 microscope (×400). For the seed coat cross-sections, four traits were investigated including the Palisade layer (or Malpighian cells), Osteosclereids, inner Parenchyma tissue, and total seed coat thickness was measured with a Zoom browser ex. Ink program (Table S1&S2; Fig. S4).

### ISSR analysis

For ISSR analysis, 31 species were planted and young leaves (0.5 g) from each accession were chosen. Total genomic DNA was extracted according to the Murray and Thomson method (1980) and the extracts from all the accessions belonging to one species were mixed. Moreover, the quality and quantity of DNA were checked by agarose gel (0.7%) electrophoresis. Of the 45 ISSR primers screened, 22 produced a higher number of reproducible bands, which were selected for ISSR analysis (Table 2). The ISSR reaction mixture (total volume = 15 ml) contained 20 ng total DNA, 1.5 mM 10x PCR buffer, 1.5 mM MgCl_2_, 0.3 mM dNTP, 2 pM of each primer, and 1 U Taq DNA polymerase. Amplifications were performed using a Bio-Rad Thermocycler (58BR 08334) programmed for 4 min at 94 °C as the initial denaturation temperature followed by 35 cycles at 94 °C for 1 min, 45 S at an appropriate annealing temperature (Table 2), 2 min at 72 °C, and 7 min at 72 °C as the last synthesis step. Amplified DNA fragments were separated in a 1.5% agarose gel at 100 W for 2 h in 1× TBE buffer (100 mM Tris–Borate, pH 8.0, 2 mM EDTA) and stained with ethidium bromide.

**Table 2:** ISSR primers information used in this study.

| Num. | Sequence (3'-5') | T<sub>a</sub> | NB | NPB | PP% | H | PIC | E | H.av | MI | D | R<sub>p</sub> |
| --- | --- | --- | --- | --- | --- | --- | --- | --- | --- | --- | --- | --- |
| 1 | (CA)8G | 52 | 14 | 11 | 0.79 | 0.494 | 0.372 | 6.097 | 0.0014 | 3.23 | 0.694 | 7.419 |
| 2 | (TC)8C | 56 | 15 | 15 | 1.00 | 0.484 | 0.367 | 6.161 | 0.0010 | 5.51 | 0.832 | 10.000 |
| 3 | (TC)8G | 54 | 14 | 12 | 0.86 | 0.472 | 0.361 | 4.581 | 0.0013 | 3.73 | 0.855 | 7.548 |
| 4 | (AC)8 G | 48 | 10 | 9 | 0.90 | 0.499 | 0.375 | 4.355 | 0.0018 | 3.04 | 0.767 | 7.032 |
| 5 | (CA)8- RT | 46 | 7 | 5 | 0.71 | 0.487 | 0.368 | 2.903 | 0.0031 | 1.31 | 0.664 | 3.161 |
| 6 | (GA)8- RT | 51 | 14 | 11 | 0.79 | 0.500 | 0.375 | 5.452 | 0.0015 | 3.26 | 0.755 | 7.097 |
| 7 | (AC)7 DBD | 50 | 12 | 10 | 0.83 | 0.475 | 0.362 | 6.742 | 0.0014 | 3.66 | 0.625 | 7.226 |
| 8 | (AG)7C | 52 | 9 | 9 | 1.00 | 0.487 | 0.368 | 3.774 | 0.0017 | 3.31 | 0.825 | 4.000 |
| 9 | (GA)8SC | 57 | 12 | 11 | 0.92 | 0.494 | 0.372 | 6.097 | 0.0014 | 3.76 | 0.694 | 7.613 |
| 10 | (AC)8C | 48 | 8 | 7 | 0.88 | 0.500 | 0.375 | 3.548 | 0.0023 | 2.31 | 0.744 | 4.258 |
| 11 | (AG)8 SG | 56 | 10 | 10 | 1.00 | 0.500 | 0.374 | 4.903 | 0.0016 | 3.74 | 0.760 | 6.387 |
| 12 | (GA)8 SG | 58 | 14 | 12 | 0.86 | 0.483 | 0.366 | 7.710 | 0.0012 | 4.42 | 0.649 | 8.065 |
| 13 | (GA)8 WT | 47 | 11 | 11 | 1.00 | 0.486 | 0.368 | 6.419 | 0.0014 | 4.05 | 0.660 | 8.129 |
| 14 | (CT)8-RG | 51 | 10 | 9 | 0.9 | 0.487 | 0.368 | 4.194 | 0.0016 | 3.68 | 0.825 | 5.548 |
| 15 | (GA)8-C | 50 | 12 | 10 | 0.83 | 0.437 | 0.342 | 6.774 | 0.0014 | 2.84 | 0.542 | 5.677 |
| 16 | (AC)8-T | 54 | 13 | 10 | 0.77 | 0.423 | 0.333 | 6.968 | 0.0014 | 2.56 | 0.515 | 5.871 |
| 17 | (GA)8-YT | 52 | 11 | 10 | 0.91 | 0.490 | 0.370 | 5.710 | 0.0016 | 3.37 | 0.675 | 7.032 |
| 18 | (GA)8-YC | 54 | 12 | 11 | 0.92 | 0.473 | 0.361 | 4.226 | 0.0014 | 3.65 | 0.853 | 6.968 |
| 19 | (AG)8-YT | 54 | 10 | 10 | 1.00 | 0.485 | 0.367 | 4.129 | 0.0016 | 3.67 | 0.830 | 6.000 |
| 20 | (GACA)4 | 50 | 12 | 9 | 0.75 | 0.498 | 0.374 | 4.774 | 0.0018 | 2.52 | 0.720 | 7.355 |
| 21 | (GA)8- RC | 51 | 12 | 11 | 0.92 | 0.497 | 0.373 | 5.935 | 0.0015 | 3.77 | 0.710 | 8.710 |
| 22 | (GACA)5 | 55 | 10 | 10 | 1.00 | 0.496 | 0.373 | 5.419 | 0.0016 | 3.73 | 0.707 | 5.742 |

T<sub>a</sub> = Annealing temperature, NB: Number of total band, NPB: Number of polymorphic band, PP%: Percentage of polymorphic band, H: Heterozygosity index, PIC: Polymorphic Information Content, E: Effective multiplex ratio, H.av: Arithmetic mean of Heterozygosity index, MI: Marker Index, D: Discriminating power, and R: Resolving power

### Statistical analysis

The fruit and seed morphological and anatomical characteristics were coded between 0 (absent) and 1 (present) to create the data matrix for computation (Table S2). For molecular analysis, only the sharp, distinct, and well-resolved fragments were scored as 1 designating present and 0 designating absent as part of a binary matrix (Table S3). Moreover, to increase the ISSR band accuracy, the fragment without any missing data was chosen. So, out of 252 bands, only 223 were used for downstream analysis. To calculate different parameters related to the primer performance, the iMEC program (https://irscope.shinyapps.io/iMEC/) was used (Amiryousefi *et al*., 2018). These parameters are Information Content (PIC), Resolving Power (Rp), Discriminating Power (Dp), expected heterozygosity (H), arithmetic mean heterozygosity (H.avp), and Effective multiplex ratio (E) (Table 2). Moreover, genetic similarity among all the species was calculated according to the Jaccard similarity index with NTSYS (ver. 2.02) (Rohlf, 1998) (Table S5) and principal component analysis (PCA) was carried out with NCSS software (2021).

To measure the genetic divergence between *Onobrychis* species, Euclidean’s genetic distances matrix was used and tree were implemented in the factoextra package in R version 3.4.2 (R Project for Statistical Computing, www.r-project.org/). Furthermore, to better show the relationships between the *Onobrychis* species, both seed morphological traits and ISSR marker data were combined. The phylogenic tree-based of joint data was carried out with Maximum Likelihood methods with 1000 bootstrap in MEGA version 6.10 for Windows (Tamura *et al*., 2013). In this study, parameters related to the genetic structure were evaluated by the POPGENE program (ver. 1.32) (Yeh *et al*., 1999). In addition, Arlequin (ver. 3.5.2.2) (Excoffier and Lischer, 2010) was used to investigate the diversity between subgroups and sections. For genetic structure and admixture detection, a Bayesian model clustering algorithm by STRUCTURE, version 2.3.4 was performed (Pritchard *et al*., 2010). For estimation of the most likely number of genetic groups (K), the output files of structure analysis were compressed into a single file then uploaded online using Structure Harvester (ver. 0.6.93) (http://taylor0.biology.ucla.edu/structure) (Earl., 2012).

## Result

### Seed and fruit Morphological Study

Seed and fruit morphological characteristics of the species are shown in Tables S1&S2. The pod was acanaceous in all the species, except in *O. pulchella*. On the other hand, the only species with short teeth on spines (component spines) was *O. crista-galli* while other species had simple spines (Table S2). Among the members of the *Onobrychis* genus studied, only two species (*O. hajastana* and *O. crista-galli*) had thorns with triangular cross-sections on the disc while other species had conical or both conical and triangular cross section spines on disc (Table S2 and Fig. S1). The marginal spine cross-sections in all the species were conical except those belonging to the *Lophobrychis* and *Onobrychis* sections. The surface of the pod in all the species was villous except for the *O. persica*, *O. pulchella*, and *O. crista-galli* while the only species with silky hair was *O. melanotricha* (Table S2). The result showed that all the species belonging to the *Hymenobrychis* section had sub-marginal veins and borders. On the other hand, only two species (*O. melanotricha* and *O. argyrea*) had no crest on fruits (Table S2&Fig. S1). Furthermore, all the species of the *Hymenobrychis* sections had fully-curved crests. The size of the legumes ranged between 1.49 mm (*O. pulchella*) and 2.91 mm (*O. arenaria*) in length, 9.74 mm (*O. pulchella*) and 1.98 mm (*O. arenaria*) in width, and 2.37 (*O. caput-galli*) and 1.125 (*O. petolemaica*) in length/width ratio. Among the members of the *Onobrychis* genus studied, only the *O. crista-galli* and species belonging to the *Hymenobrychis* section were found to have two seeds in fruit while the other species contained a single seed in fruit (Table S2). The seed shapes were distinguished as Trapeziform, Orbiculate, Rhombic, Reniform, and Ovate. However, the general shape of the seeds in the four sections was reniform (Table S2).

Based on 40 species of the *Onobrychis* genus, Yildiz *et al*. (1999) suggested three morphology types for pods (*O. radiata* type, *O. beata* type, and *O. ornata* type) in the *Onobrychis* genus. However, the species in this study, especially those having the same pod shapes as the *O. beata* type were varied in legume shape, spines density, crest length, and spine type. Hence, based on our results, recognition of another pod type seems to be necessary for the *Onobrychis* genus; we propose the *O. crista galli* type to be recognized in addition to the three already recognized fruit types for this genus (Table S1&S2 and Fig. S1). So, a key is now provided for distinguishing between fruit (legume) types. Type 1 (*O. beata* type) involved the simple spine type, both triangular and conical transverse spine sections on margin and disk, curved crest as a half circle with a short crest, and one seed. This was observed in all the species of the *Onobrychis* section and in some of the *Lophobrychis* sections such as *O. aequidentata*, *O. pulchella*, and *O. caput-galli* (Table S2& Fig. S1). Although *O. pulchella* was the only species without hair and spines, the general shape was the same as type one. It may, therefore, be categorized with type one species (Table S2 and Fig. S1). Type 2 (*O. radiata* type) consisted of the simple spine type, conical surface spine section, conical marginal spine section, a fully-curved crest with a long crest, marginal veins, and 2 ovulate ovate. All the species of the *Hymenobrychis* section were found to have this fruit type. Type 3 (*O. crista galli* type) included the compound spine type with a high surface spine density; long spines; glabrous, curved crest as a half circle with a short crest, and 2 ovulate ovates. Based on our results, this type is only observed in *O. crista-galli*. Type 4 (*O. ornate* type) contained the one with the simple spine type with a conical surface spine section, a conical marginal spine section without a crest but with marginal veins and borders. This was observed in the species of the *Heliobrychis* section (Table S2).

### Seed micro-morphological study

#### Light microscopy study

The general anatomy of the *Onobrychis* seed coat was found to be similar to that described by Cenci et al. (2000) who reported the outer layer represented by radially elongated cells to form a palisade-like (Malpighian cells) layer that constitutes the outer epidermis. Below the Malpighian cells, there are bone-shaped and thick-walled cells, termed ‘osteosclereids’, which underlie the inner tissue with thin-walled parenchyma cells which is followed by the cuticle layer (Fig. S4). In this study, *O. pulchella* showed the highest values for the Malpighian cells while *O. melanotricha* had the lowest values for the osteosclerosis layer. Significant differences were also observed in the total thickness of the parenchyma layer, with *O. michauxii* having the highest (0.88 mm) and *O. pallasii* having the lowest value (0.024 mm). The total seed coat thickness ranged between 0.142 mm and 0.052 mm in *O. pulchella* and *O. melanotricha*, respectively.

#### Scanning electron microscopy study (SEM)

The seed coat sculpture patterns were classified into six types: Reticulate (having a network wrinkle), Undulate (having wavy wrinkles), Druse (having aggregation of irregular wrinkles), Convolutus (having tortuous wrinkles), Flat (having flat wrinkles), and Stellate (Asteroid elevations) (Table S1 and Fig. S2). The majority of the studied species showed the reticulate (14 species) followed by Undulate (5 species), Convolutus (8 species), flat (2 species) and the remaining two types each represented by only one single species (Table S2 and Fig. S2). Cluster analysis of seed & fruit morphological data

Based on 75 seed morphological traits, the species were divided into two main groups (Fig. 1A). Group A consisted of all the species from the *Onobrychis*, *Lophobrychis* sections except *O. aequidentata* while the second group contains all the species from the *Hymenobrychis* and *Heliobrychis* sections along with *O. pulchella* (Fig. 1A). As a complementary test in grouping species with minimal data loss, we performed principal components analysis in which the first three principal components covered 41.79% of the cumulative variation (Table S4). Two-dimensional visual illustrations of the first three PCA are shown in Figs. 2A & S5. Based on the first two components, two district groups were found. The first group contains all the *Onobrychis* subgenera’s species, despite the small distance of the *O. pulchella* due to first PC. The second group contains all *Sisyrosema* subgenus species in which almost all the species related to the section *Heliobrychis* were separated from the section *Hymenobrychis* according to the second PC (Fig.2A).

**Fig 1.**
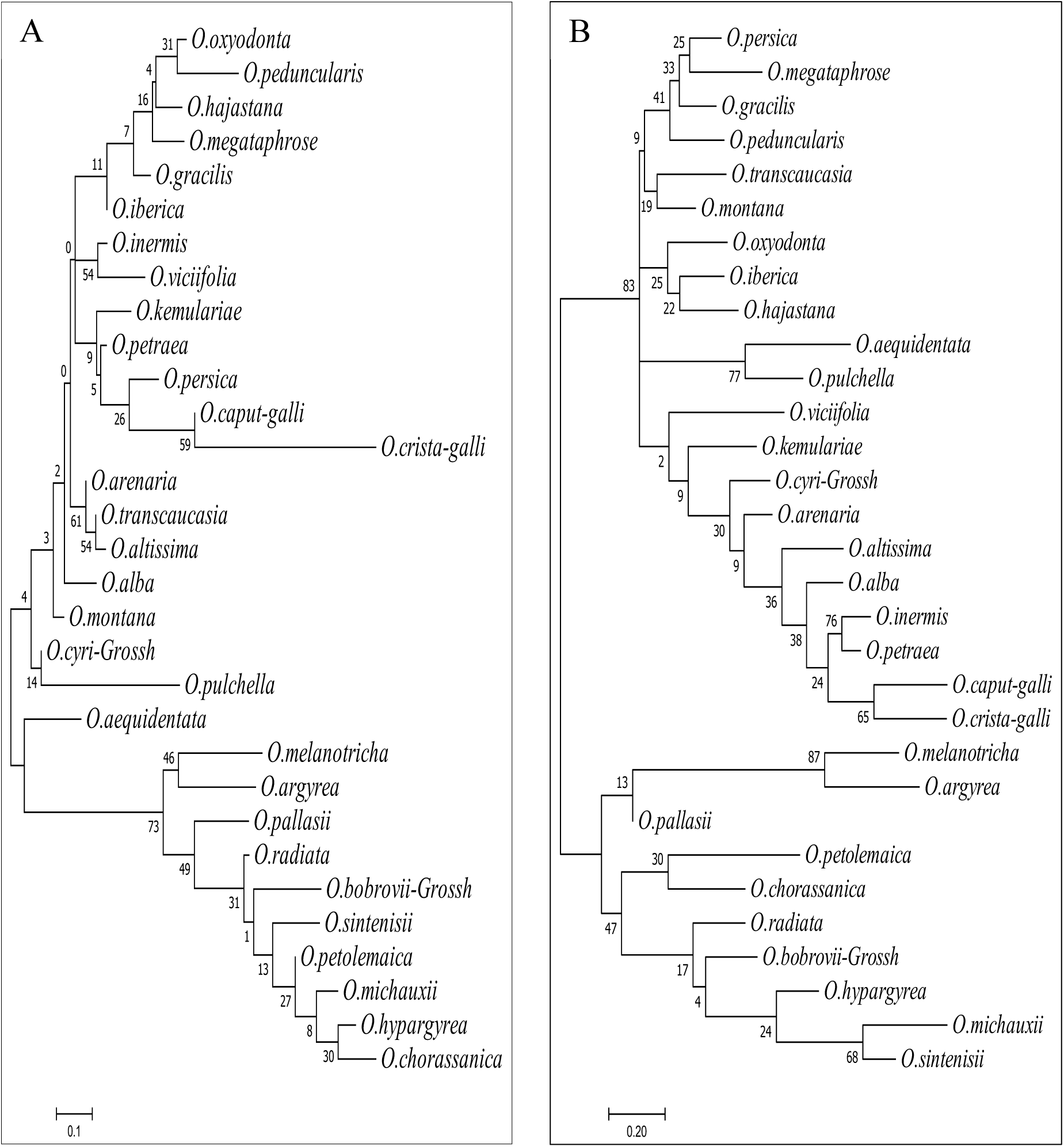
Association between *Onobrychis* species reveled by complete linkage model and Euclidean’s genetic distance based on seed and fruit macro & micro-morphological traits (A) and ISSR marker (B).

**Fig 2.**
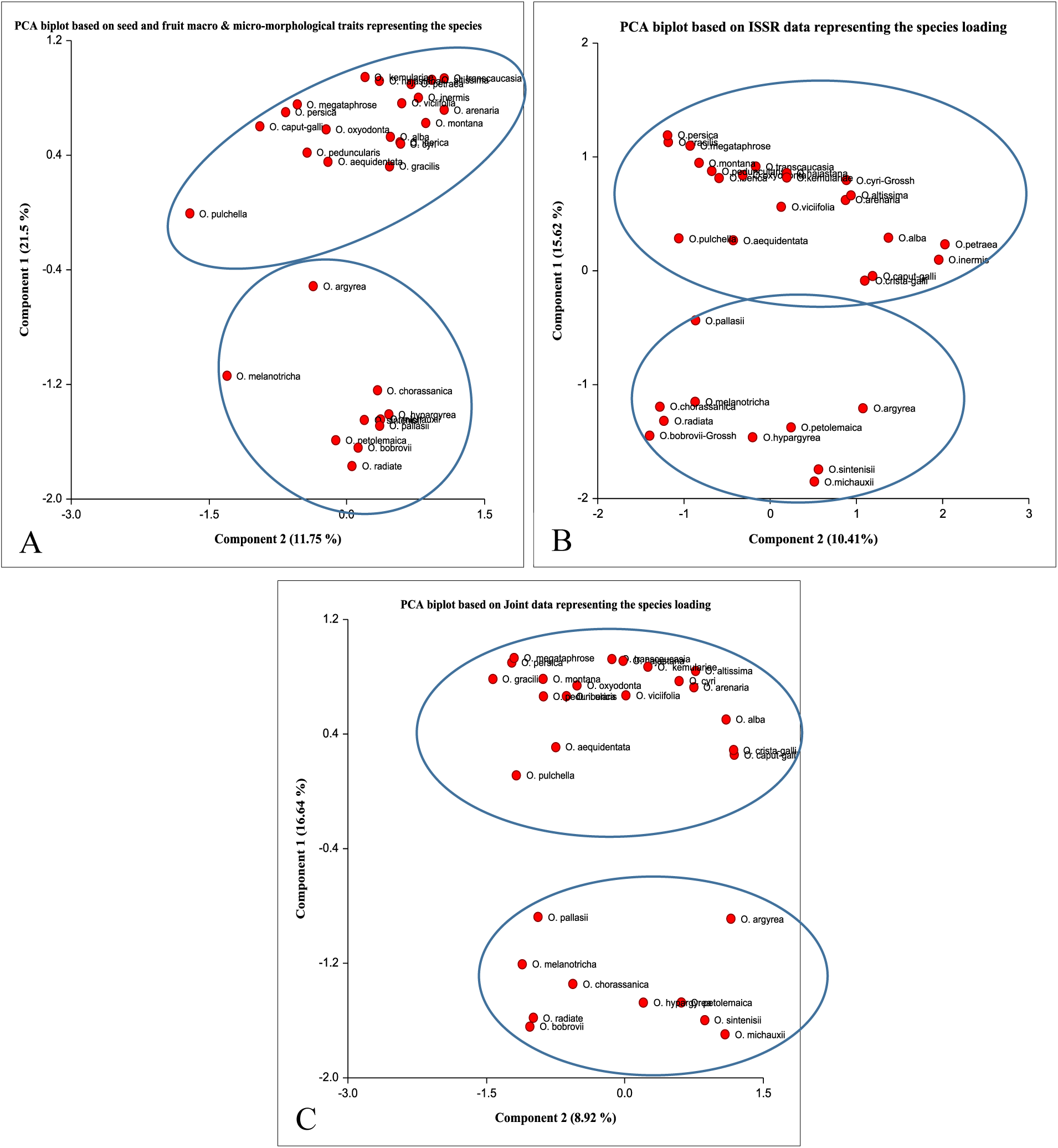
PCA (correlation’s measure) case scores using NCSS program. A: Seed and Fruit macro & micro-morphological traits, B: ISSR marker, and C: Joint data.

### ISSR analysis

#### Marker efficiency

The 22 primers selected amplified 252 bands (Table S3), of which 223 (88.5%) were polymorphic among 31 *Onobrychis* species. The amplified DNA ranged from 150 to 1400 bp. The maximum and minimum numbers of bands were observed by the primers (TC)_8_-C and (CA)_8_-RT, respectively. The percentage of polymorphic bands ranged from 75% to 100% (Table 2). The heterozygosity (H) varied from 0.423 (ISSR 16) to 0.5 for ISSR 06&10&11 with a mean of 0.484 per primer. The polymorphism information content (PIC) value ranged from 0.33% ((AC)8-T) to 0.38% ((GA)8-RT) with an average of 0.37%. Moreover, the marker index (MI) with an average value of 0.008 ranged from 0.006 to 0.01 (Table 2). For resolving power index (Rp), the lowest and highest values belonged to (CA)8-RT (3.16) and (TC)8C (10), respectively (Table 2). The arithmetic means of heterozygosity (H.av) ranged between 0.001 and 0.0031 with the mean of 0.0016 per primer. To determine the judicious profundity of primer, we calculate discriminative power (D) with a mean index of 0.723 and extend it from 0.515 ((AC)8-T) to 0.855 ((TC)8G). The effective multiplex ratio (E), which is a conditional factor on the magnitude of primer polymorphism, is spanned from 2.9 ((CA)8-RT) to 7.71 ((GA)8 SG) with an average of 5.312 per primer.

#### ISSR based Genetic similarity (GS)

Genetic distances and similarity coefficients for different species (Jaccard methods) and sections (Ni & Li method) are calculated and are shown in the Table S5 and Table S6, respectively. The findings of the present study revealed the significant genetic differentiation among the *Onobrychis* species which ranged from 0.199 (*O. bobrovii* vs. *O. megataphrose*) to 0.833 (*O. petraea* vas *O. inermis*) (Table S5). Moreover, from a sectional classification point of view, the highest similarity was observed between *Onobrychis* and *Lophobrychis* sections (0.87) while the lowest similarity was calculated for *Lophobrychis* and *Heliobrychis* (0.61) (Table S6).

#### ISSR Cluster analysis

The cluster analysis based on ISSR marker was performed for the 31 species as shown in Fig. 1B. The species were divided into two main groups in which all species belong to *Onobrychis* and *Sisyrosema* subgenus grouped in the first and second clades, respectively (Fig. 1B); however, in the first group, all species belonging to the *Onobrychis* subgenera are mixed, while in the second group *Hymenobrychis* and *Heliobrychis* sections are mostly separated (Fig. 1B).

For more validation, the principal component analysis was run. The first three principal components cumulatively explained 32.36 % of all the variation (Table S4). Like the cluster analysis, we observed complete discrimination of subgenus. However, unlike cluster analysis, *Sisyrosema* sectional classification was not observed, and also, based on the first component *O. pallasii,* few were grouped far from their other subgenus member (Fig. 2B and Fig. S5).

#### Genetic Structure

Estimates of population parameters in each subgroup are shown in Table 3. The results showed that there was a slight difference between the calculated parameters for both subgenera. However, the calculation of the parameter for each section showed that the greatest variation was related to *Onobrychis* and *Hymenobrychis* sections equally (Na: 1.8, Ne: 1.5, H: 0.3, and I: 0.44) while the lowest variation belongs to *Heliobrychis* section (Na: 1.34, Ne: 1.24, H: 0.14, and I: 0.21).

**Table 3:** genetic parameter of *Onobrychis* subgenera and sections

| Table 3 : genetic parameter of *Onobrychis* subgenera and sections |  |  |  |  |  |  |  |  |
| --- | --- | --- | --- | --- | --- | --- | --- | --- |
| subgenus | section | No. of species | Na | Ne | H | I | Number of polymer | PPB% |
| *Onobrychis* |  | 21 | 1.84 | 1.54 | 0.31 | 0.47 | 189 | 84.75 |
|  | *Onobrychis* | 17 | 1.8 | 1.51 | 0.3 | 0.44 | 178 | 79.82 |
|  | *Lophobrychis* | 4 | 1.68 | 1.47 | 0.27 | 0.4 | 153 | 68.61 |
| *Sisyrosema* |  | 10 | 1.86 | 1.57 | 0.33 | 0.49 | 192 | 86.1 |
|  | *Heliobrychis* | 2 | 1.34 | 1.24 | 0.14 | 0.21 | 77 | 34.53 |
|  | *Hymenobrychis* | 8 | 1.81 | 1.5 | 0.3 | 0.44 | 182 | 81.61 |
| Na: Observed number of alleles , Ne: Effective number of alleles ,H: gene diversity , I: Shannon's index , PPB%: The percentage of polymorphic loci |  |  |  |  |  |  |  |  |

In addition, to study the structure of the existing diversity in more detail, molecular analysis of variance (AMOVA) was used (Table 4). Based on the AMOVA result, 7.8% of the total variation was related to the subgenus difference, while 24.51% of the diversity was related to the differences between the sections in each subgenus and 68.41% of the variations were related to the differences between the species in each section.

**Table 4.** : The Analysis of Molecular Variance(AMOVA) among 31 *Onobrychis*

| Table 4 : The Analysis of Molecular Variance(AMOVA) among 31 *Onobrychis* species |  |  |  |  |  |
| --- | --- | --- | --- | --- | --- |
| source of variation | Df | sum of squares | variance components | percentage of variation | p-value |
| among subgenus | 1 | 416.42 | 3.87 | 7.08 | <0.001*** |
| among section within subgenus | 2 | 355.79 | 13.39 | 24.51 | 0.0001*** |
| among species within sections | 59 | 2205.56 | 37.38 | 68.41 | 0.0001*** |
| Total | 62 | 2977.77 | 54.64 |  |  |

For evaluation of population structure and assigning species to specific populations based on ISSR marker, Structure version 2.2.3 was used based on a Bayesian model. The structure analysis was initially performed based on the maximum number of (K = 1 to 7), as the original population order and the most probable value of population were calculated to the maximum peak at ΔK = 4 (K value =3.25; Lnprob (K) = -3669.8). The rate of change of the likelihood distribution (mean), the absolute value of the 2^nd^ order of the change of the likelihood distribution (mean) for ΔK = 4 are shown in Fig. 3. Based on best K = 4, it was determined that all the evaluated species could be positioned into four major groups visualized with four distinct colors of blue, green, yellow, and red. Each individual is represented by one vertical bar plot and the labels below the bar plots are the corresponding numbers for each species in Table 1. More than one color in each bar plot reveals the genetic complexity of that individual that, in this case, the species belongs to a group that has the largest color width of that cluster. The estimated mean F_st_ value for the species under clusters 1 and 2 was 0.2412 and 0.279, respectively while this value for clusters 3 and 4 was 0.515 and 0.823, respectively (Table S7). Due to the membership likelihood, described by We *et al*., (49), some of the species showed unique structure. Based on Q > 0.60 as the purity standard, in the red zone 7 species, almost 5 species in the blue zone, and 3 species in the yellow zone shows some level of purity and the rest of the species were noted as admixture units based on Q < 0.60 (Fig. 3B).

**Fig 3.**
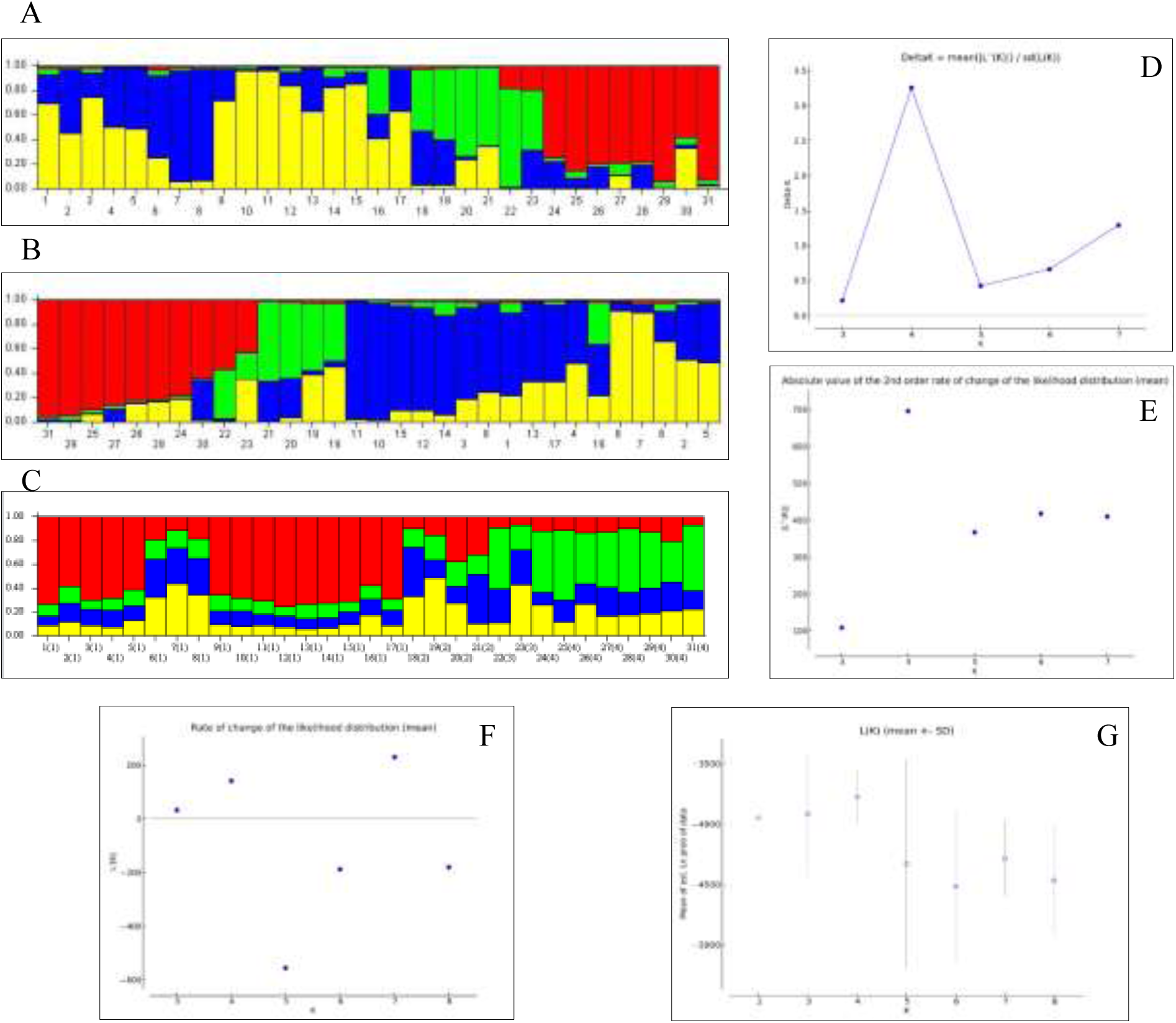
Structure harvester and Delta K value Bayesian model based assessment of population structure for 31 *Onobrychis* species based on ISSR markers. A: Bar Plot based on original population order, B: Bar plot based on estimated membership coefficients values, C: Bar Plot based on species ID (population) order, D: ΔK = mean (|L”(K)|)/sd(L(K)) and ΔK = 4 indicates the best K value, E: absolute value of the 2^nd^ order rate of change of the likelihood distribution (mean), F: rate of change of the likelihood distribution (mean), and G: mean of estimated Ln probability.

#### Seed and fruit morphological and ISSR combination analysis

To better discriminate between the *Onobrychis* species, both seed morphological traits and ISSR marker data were combined. So, based on joint data, Jaccard’s Genetic similarity coefficients (GS) between the studied species were calculated and were in the range between 0.185 (*O. melanotricha* vs *O. persica*) and 0.806 (*O. inermis* vs *O. petraea*) (Table S5). The GS mean value for each section was 0.5, 0.43, 0.52, and 0.49 for *Onobrychis*, *Lophobrychis*, *Heliobrychis*, and *Hymenobrychis,* respectively (Table S5).

The hierarchical clustering analysis based on Joint data is shown in Fig. 4. The dendrogram illustrated that all the 31 species were clustered into two main groups. Group A consisted of all *Onobrychis* subgenus species in which *O. arenaria* was classified in a separate subgroup. Furthermore, all the *Hymenobrychis* and *Heliobrychis* section species were clustered into the second group with complete separation (Fig. 4).

**Fig 4.**
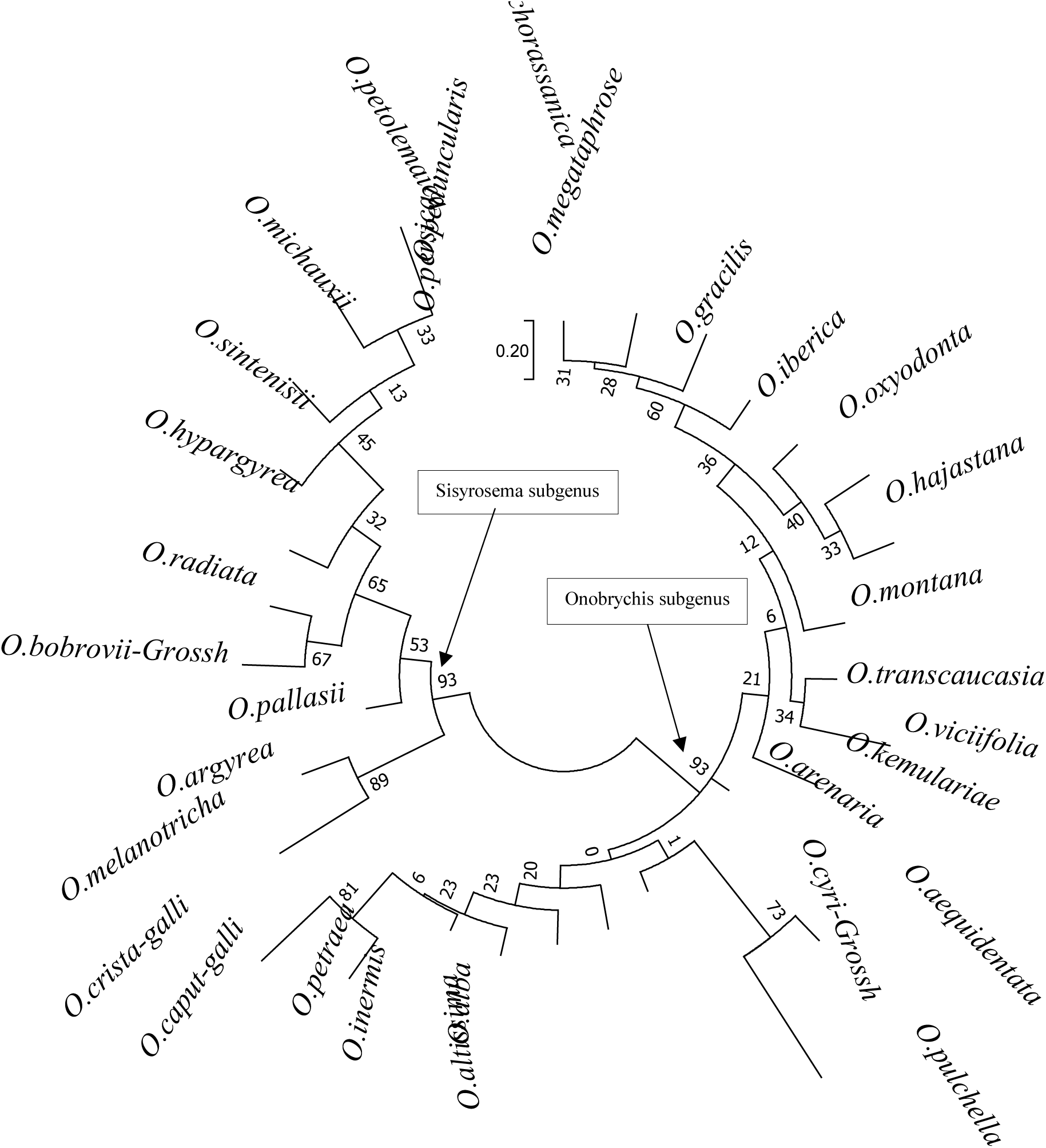
The Maximum Likelihood phylogenetic tree illustrating the genetic relationship among *Onobrychis* species based on 1000 bootstrap based on Maximum Likelihood model.

To lead the clustering investigation, principal component analysis was used. The distribution of eigenvalues, percentage of genetic variation, and cumulative percentage of genetic variation were displayed in Table S4. The PCA analysis revealed that the share of the first three PC is 41.77 % (PC1 = 18.85%, PC2 =13.22% and PC3 = 9.7%) of total variation. In the case of PCA analysis from the first principal component, the highest value was for *O. megataphrose* and *O. transcaucasia* (0.93 and 0.92 respectively) while the least value (-1.96) was found for *O. michauxii* which has the least contribution to total diversity (Table S4). The first two PC bi-plots showed the complete separation of the subgenus classification (Fig. 2C). However, as far as the sections are concerned, this discrimination has not happened.

### Discussion

#### Seed and pod morphological characteristics

Legume morphological characteristics, seed-coat sculpturing, and seed-coat anatomy have been shown to provide valuable characters in the delimitation of selected species of the *Onobrychis* genus. Norri and colleagues (2014) showed that morpho-biometerical characters are useful in distinguishing among species and pod characters are the most valuable issue for separation of the *Onobrychis* genus. A great variation in the morphological characteristics in this study showed they could be useful tools for classifying and identifying species in the *Onobrychis* genus. However, none of the fruit morphological traits are specific to one section or subgenus. For example, we can refer to the fully carve crest in the fruit, which can be considered as an indicator for the *Hymenobrychis* section, nonetheless, this indicator did not exist in the *O. pallasii*. These traits in combination with each other might be used as an identification key for the corresponding section or subgenre. Therefore, in addition to the different types of fruit that were suggested by Yildiz *et al*. (1999), we also introduced a new fruit type called *O. crista* galli. Apart from the differences between new fruit type with *O. beata* type mentioned earlier (legume shape, spines density, crest length, and spin type), a fundamental difference between these two types is the number of seeds in the fruit, which for *O. beata* type this number is one, while the newly introduced type has ovary 2-ovulate (Table S2). Due to the characteristics related to fruit, a very interesting result is the high variability in the *Lophobrychis* section. Two species in this section had unique characteristics. The first species was *O. pulchella*, which generally has fruit like *O. beata* type, but it did not have any thorns on the fruit or long crest length (Table S2 and Fig. S1). The second species was *O. crista* galli, which was the only species in this study that had a compound spin and unlike all species of *Lophobrychis* section, had the ovary with 2-ovulate (Table S2 and Fig. S1).

For the seed transverse section, no significant variation was observed in the osteosclerotic layer in any of the species while the difference in seed coat thickness was mainly due to variations in the length of the Malpighian cells and the inner parenchyma tissues (data not shown). Contrary to Cenci *et al*. (2000), who reported differences in the shape of Malpighian cells between the perennial and annual species, we did not observe any significant differences between them (Table S2).

For seed coat patterns, the variation observed was generally species-specific rather than genus-specific and almost invariant across the different populations of a given species (Koul *et al*., 2000 and Moazzeni *et al*., 2007). Our results revealed six main types of seed-coat patterns among the investigated species (Table S2 and Fig. S3). Although it seems that the seed-coat patterns observed in this study were variable, it was not sections or subgenus specific. Therefore, it cannot be used as a unique trait to classify species of this genus.

Another interesting result in our study is the differentiation between annual and perennial species in *Onobrychis* genus. Based on Flora Iranica (Rechinger, 1969), Flora of Turkey (Davis *et al*. 1988), and Flora Europaea (Ball, 1968), all the species belonging to the *Lophobrychis* section are annual while all the other sections have perennial species. However, we did not find any differences among the annual and perennial species, neither by seed anatomical and fruit morphological traits nor by ISSR markers (Fig. 1). Similar results have been reported by Pavlova and Monova (2000) who found no differences in pollen morphology among the annual and perennial species in the *Onobrychis* genus.

#### ISSR marker

It has been well documented that ISSR markers are highly polymorphic and are useful tools for studying genetic diversity, phylogeny, gene tagging, genome mapping and evolutionary biology (Pradeep *et al*., 2002). Our ISSR marker results represented a good level of polymorphism suggesting the efficacy of these markers for the evaluation of the phylogenic relationship among the *Onobrychis* species. There are many factors to check the efficacy of primers, some of which have great importance. Two major indexes for the informativeness of genomic marker polymorphism are expected heterozygosity (H) and the polymorphic information content (PIC). PIC and H measures the ability of a marker to detect polymorphisms, and is important in selecting markers for genetic studies (Khan *et al*., 2021). Considering, the range of H and PIC value for a dominant marker (0 for monomorphic primer to 0.5 (for primer with multiple alleles in an identical frequency), the value for these two indexes in studied population was more than average, which indicated the advanced discriminatory capacity of the marker system. The MI index, in addition to the benefits of PIC, also considers the total number of bands and the polymorphic ratio which shows the potential of each primer to produce more bands. Moreover, Rp measures the ability of primers to recognize and differentiate individuals. So, all the represented values for these indexes showed the high polymorphism and efficiency of the used primer for genetic discrimination analysis on this genus (Table 2). Overall, the greater RP and MI indices refer to the greater efficacy of the respective primers (Zarei *et al*., 2021). Therefore, the most effective primers are (TC)8C (Rp= 10, MI = 5.51), (GA)8-RC (Rp = 8.7, MI= 3.77), (GA)8 WT (Rp=8.12, MI= 4.05) and (GA)8 SG (Rp= 8, MI = 4.42)

#### Genetic structure based on ISSR marker

The population structure of *Onobrychis* genus assessed by Bayesian admixture analysis indicated 4 clusters (Fig. 4). Based on Q > 0.60, out of 31 *Onobrychis* species 15 were almost pure and the other species were highly complex, indicating these species were genetically admixture. Interestingly all the species belonging to *Lophobrychis* section (number 18, 19, 20 and 21) have different color in each bar plot which reveals their genetic complexity. Moreover, the bar related to *O. viciifolia* had three colors (red, blue and yellow) in almost equal proportions, which indicates the genetic complexity of this species too. Perhaps, the cross-pollinated nature of cultivated sainfoin (*O. viciifolia*) as well as vast diversity regain make this species so complex. For better understanding of differentiation among studied population, fixation index (Fst) was calculated. Due to Hartl and Clark (1997), a standard scale of fixation index categorized in: little genetic difference (< 0.05), moderate genetic difference (0.05–0.15), great genetic difference (0.15–0.25) and very great genetic difference (> 0.25). The estimated Fst suggested the species under cluster 1 (Fst = 0.241) showed little genetic difference while the species under cluster 2 (Fst = 0.279), 3 (Fst = 0.515) and 4 (Fst = 0.823) showed very great genetic difference. Over all, these Fst values indicated high level of genetic differentiation with greater genetic distance between clusters.

#### Sectional and Subgenus classification

In traditional classification, characterizing and assigning species to sections or sub-genera seem to have often been based on morphological traits such as flower and leaf characteristic or seeds and fruits related traits. So, some previous researchers, such as Yildiz and colleagues (1999), by using seeds and fruits morphology claim that no evidence can be found in support of subgenera classification while sectional discrimination was supported. On the contrary, some phylogenic studies based on molecular markers, had completely opposite results in the supporting subgenus classification and no verification of sectional discrimination, especially in the subgenus *Onobrychis*. (Ahangarian *et al*., 2007; Safaei Chaei Kar *et al*., 2012) So, considering the morphological traits of this study, even though the key was provided to identify the fruits and seeds of each section and species, not only the subgenus, but also the sectional discrimination for *Onobrychis* subgenus was not confirmed. These results show that usage of morphological characters alone in phylogenetic analysis was restricted. Such restriction happened because of the open pollination with a high allogamy rate (Avci *et al*., 2016), along with wide geographical dispersion (Zarrabian *et al*., 2013) of the *Onobrychis* taxa. Therefore, the morphological characters are more affected by environmental factors. Contrary to the morphological traits, most of the molecular data supported the subgenera classification, however, they did not support the sectional discrimination in *Onobrychis* subgenus (Ahangarian *et al*., 2007; Chaei Kar *et al*., 2014 and Duan *et al*., 2015). So, to increase the accuracy of finding relationships between species, especially relationships between sections and subgenus, joint data was used and downstream analysis was conducted. Our study with sufficient species sampling for each subgenre suggests that *Sisyrosema* is the only monophyletic subgenera. And also, contrary to *Onobrychis* subgenre, *Sisyrosema* was divided according to its traditional sectional classification. Such classification may not have occurred by principal component analysis for joint data, but at least two subgenres were clearly separated from each other (Fig. 2C). Other researchers have used seed storage protein (Arslan and Ertugrul, 2010) and sequence data of ITS, matK, trnL-F and psbA-trnH (Duan *et al*., 2015), suggesting that the subgenus *Sisyrosema* might be derived from the subgenus *Onobrychis*. In the nrDNA ITS-based phylogenic studies (Ahangarian *et al*., 2007; Safaei Chaei Kar *et al*., 2012) the non-monophyly of subgenus *Onobrychis* was also indicated, but subgenus *Sisyrosema* was retained as monophyletic. Also, Hayot-Carbonero and colleagues (2012) supported the paraphyly of *Onobrychis* subgenus. They suggested, not only from a molecular point of view, but also from morphological perspective, all species from the *Onobrychis* section are very similar to *O. viciifolia*. They also found that *O. petraeais* is more closely related to the *Lophobrychis* section, which our result completely agrees with (Table S4B).

#### *Lophobrychis* section as a heterogeneous unit

Generally, the species were positioned more closely in the generated phylogenetic tree, suggesting that they were genetically more similar or even have identical genes. On the other hand, species possessing the distant group suggest that they were genetically dissimilar, even though they come from the same origin (Khan *et al*., 2021). Such a trend has not been observed for the species related to *Lophobrychis* section. In this study, no independent clade containing all species of the *Lophobrychis* section was found, neither through molecular markers nor with morphological traits. Therefore, in contrast to the traditional taxonomic classification, species belonging to the *Lophobrychis* section were not coherently classified in one group (Fig.1 and Fig. 4), indicating the *Lophobrychis* section not only has a comparatively highly derived organization, but also can be considered as a heterogeneous unit in the *Onobrychis* genus (Abou-el-Enain, 2002; 2004). Although a previous study (Safaei Chaei Kar *et al*., 2014) showed *Onobrychis* and *Lophobrychis* sections were separated, the high similarity found in this study between *Lophobrychis* and other sections, especially the *Onobrychis* (Fig. S6 and Table S5), gives rise to the probability that the concept of section applied to *Lophobrychis* is meaningless or unrealistic. Bandara *et al*. (2013), based on ITS sequences, supported the sectional treatment and may require some changes. They assumed that, in order to form a monophyletic clade corresponding to a section, part of *Lophobrychis*, *Dendrobrychis* and the subsection *Macropterae* of *Onobrychis* section should be kept together. Hayot-Carbonero *et al*. (2012) declared the *Lophobrychis* and *Onobrychis* sections were less clearly distinct. By looking at similarity matrix based on the joint data (Table S5), O. *caput-galli* had a higher similarity to the species belonging to *Onobrychis* section rather than the rest of *Lophobrychis* species. Elena (2006) reported that *O. caput-galli* was cyto-taxonomically closer to the section *Onobrychis* rather than other species of its own section. Hence, it seems that this species could be the connecting link between the *Onobrychis* and *Lophobrychis* sections of this genus. Overall, there is a general difficulty in discrimination of the *Lophobrychis* and *Onobrychis* sections. However, it is possible that these two sections have a relatively recent common ancestor (Hayot-Carbonero *et al*., 2012).

## Conclusion

Investigations of seed macro- and micro-morphology of the genus *Onobrychis* showed that it could be a useful tool (especially macro-morphological traits) for differentiating species. Hence, these data alone are insufficient for phylogenic analysis. Such conflict may be due to the fact that species were assigned to different sections based on only one or few morphological characters and some characters may have converged in the course of evolutionary history (Duan *et al*., 2015). The results obtained from this study support the subgenus (*Onobrychis* and *Sisyrosema*) classification while the monophyly of *Onobrychis* subgenus was not supported. Moreover, it seems that the section *Lophobrychis* has a comparatively highly derived organization and can be considered as a heterogeneous unit in the *Onobrychis* genus. Our results also showed that combination of seed macro- and micro-morphological traits with ISSR data provide complementary information for the *Onobrychis* genus classification. However, more study with larger number of molecular markers will be needed to obtain an appropriate phylogenetic classification.

## Supplementary data

The following supplementary data are available at *JXB* online.

Fig. S1. : Fruit shape of different *Onobrychis* species.

Fig. S2: Transverse sections of *Onobrychis* species seed coats.

Fig. S3. Technical shape which discripe the seed surface pattern

Fig. S4: Scanning electron microscopy photographs of seed coat

Fig. S5: Two-dimensional representation of genetic relationships based on principal components analysis

Table S1. Seed and fruit macro and micro morphological characters.

Table S2: Data matrix of the fruit and seed characteristics of *Onobrychis* species in this study.

Table S3: ISSR Data matrix of species in this study.

Table S4 A: Jaccard’s Similarity matrix between the 31 *Onobrychis* species.

TableS5: Nei’s genetic identity and genetic distance for each sections based on ISSR.

Table S6: The eigenvalues, % of variation and cumulative % of variation of 30 PCs and PCoA

Table S7: species contribution in each component in different experiment

Table S8: Inferred ancestry of species and Fst value for each cluster

## Acknowledgements

I would like to thank Leibniz Institute of Plant Genetics and Crop Plant Research (IPK), and United States Department of Agriculture (USDA) for providing plant material

## Author contributions

MZ designed, analyzed and preformed the experiments; MZ, AND, MHE, MMM wrote the paper.

## Data availability

The raw data used in this study are available from the corresponding author upon request.

---

**Figure S1:**
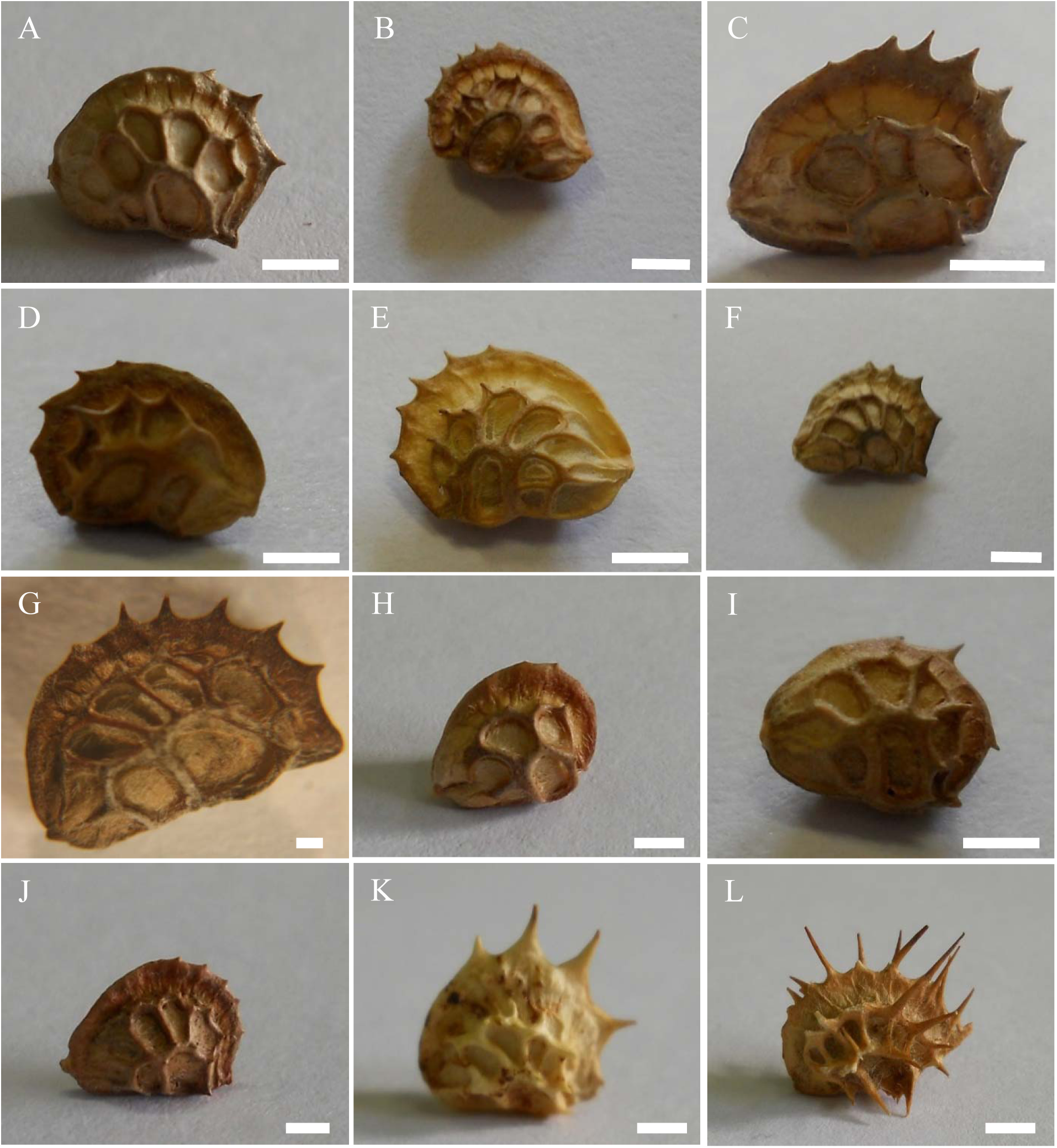
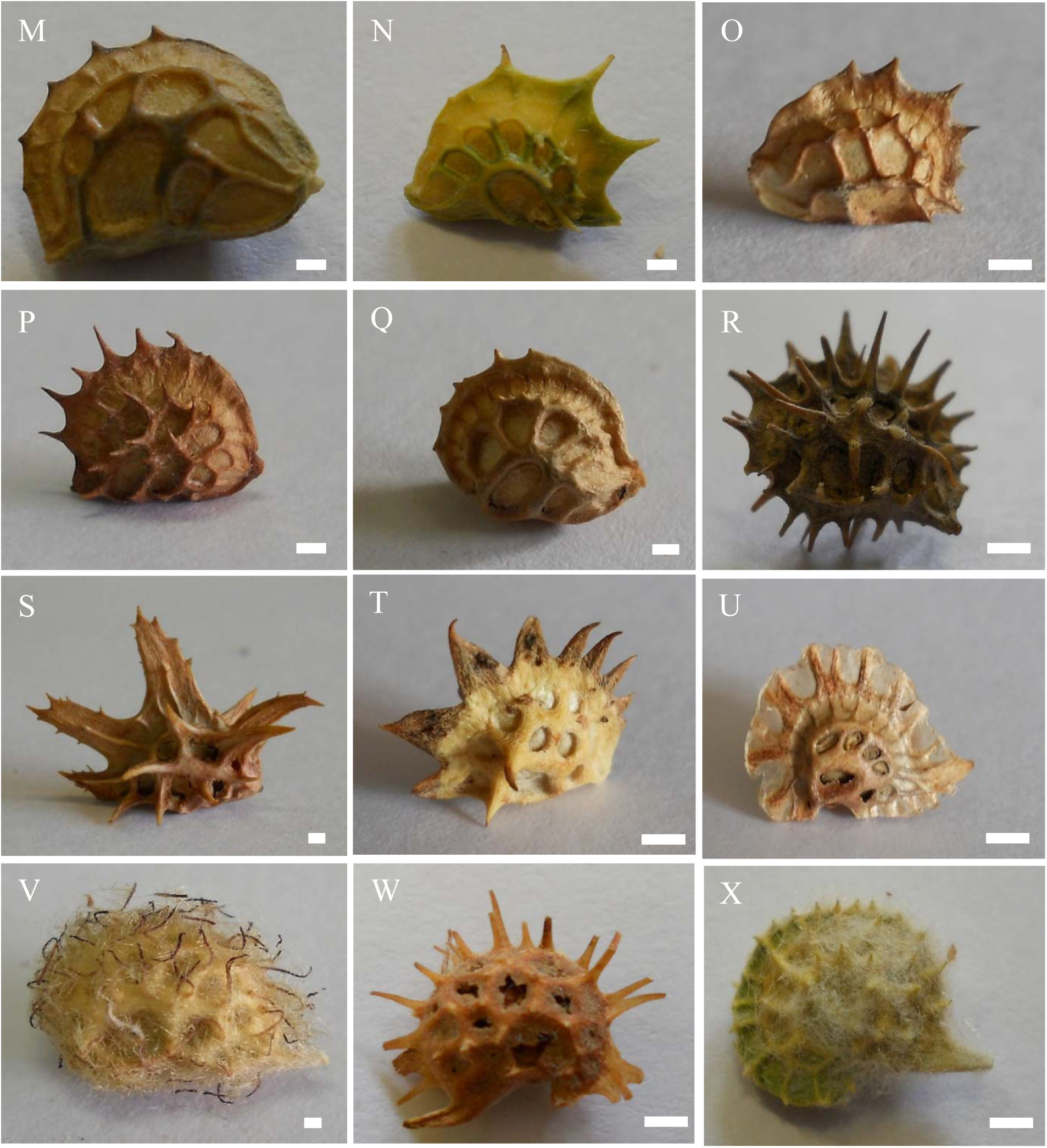
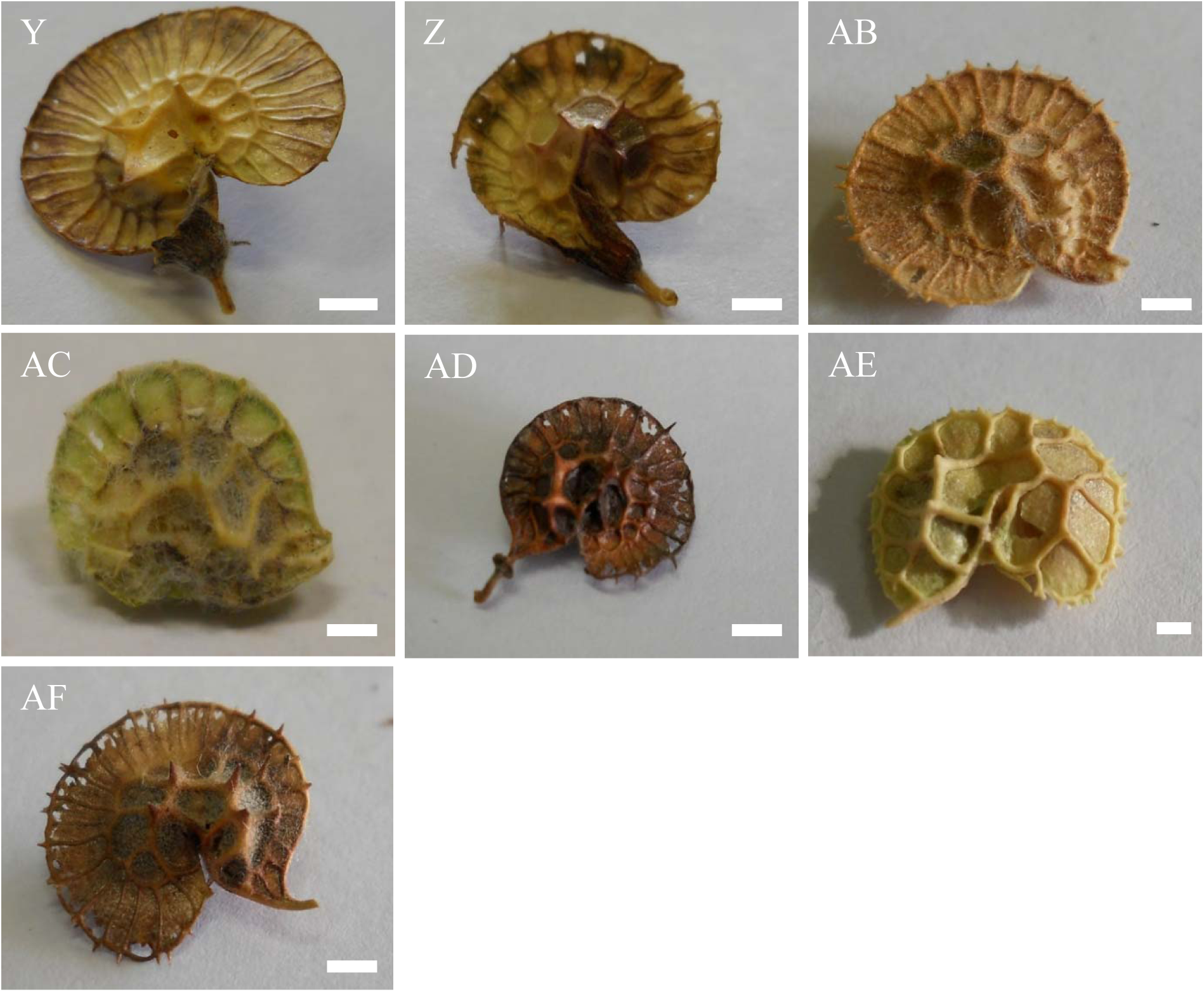
Fruit shape of different *Onobrychis* species. A: *O. transcaucasia*, B: *O. arenaria*, C: *O. iberica*, D: *O. cyri-*Grossh, E: *O. altissima*, F: *O. alba*, G: *O. inermis*, H: *O. petraea,* I: *O. oxyodonta*, J: *O. gracilis*, K: *O. persica*, L: *O. peduncularis* (Scale bar: 1 mm) M: *O. hajastana*, N: *O. megataphrose*, O: *O. montana*, P: *O. viciifolia,* Q: *O. kemulariae*, R: *O. caput-galli*, S: *O. crista-galli*, T: *O. aequidentata*, U: *O. pulchella*, V: *O. melanotricha*, W: *O.argyrea*, and X: O. ptolemaica (Scale bar: 1 mm). Y: *O. hypargyrea*, Z: *O. michauxii*, AB: *O. sintenisii*, AC: *O. chorassanica*, AD: *O. bobrovii-Grossh*, AE: *O. pallasii*, and AF: *O. radiate* (Scale bar: 1 mm).

**Figure S2:**
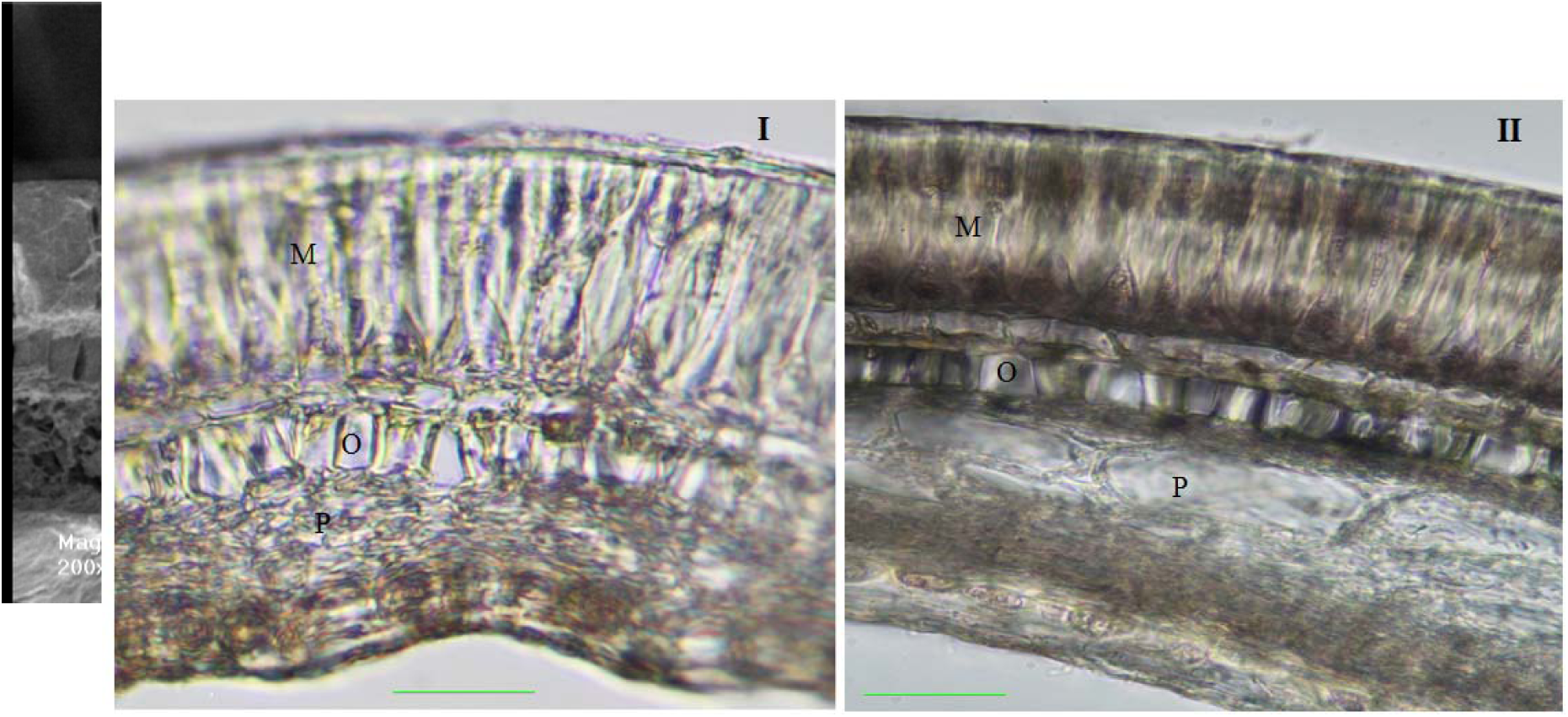
Transverse sections of *Onobrychis* species seed coats: I) *O. caput-galli,* II)_II_*O*_I_*. altissima* and III) scanning electron microscope (SEM) cross section of *O. transcaucasia*, (C= Cuticle layer; M = Malpighian cells; O= Osteosclereids; P= Inner parenchyma tissue) (Scale bar = 40 µm for Nikon ECLIPSE E600 microscope and 100 µm for SEM).

**Figure S3.**
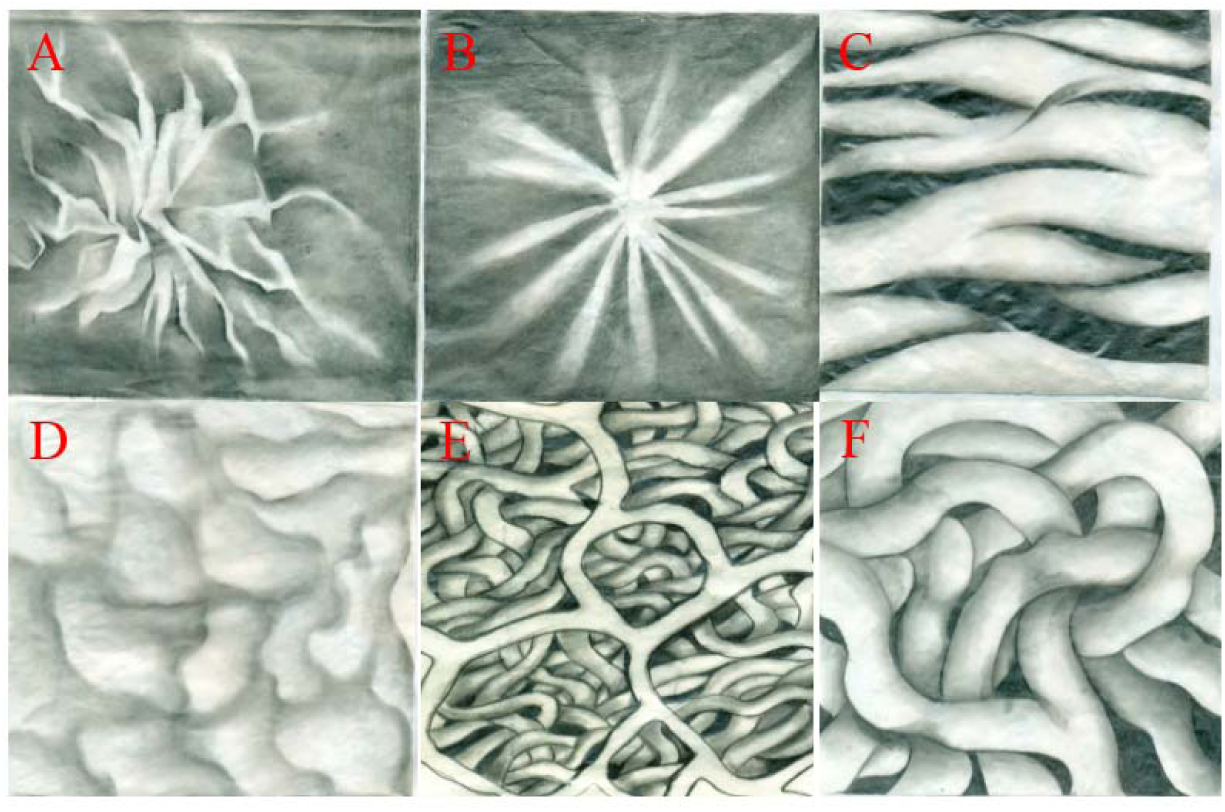
Technical shape which discripe the seed surface pattern : A) Druse, B) Stellate, C) Undulate, D) Flat, E) Reticulate, and F) Convolutus.

**Figure S4:**
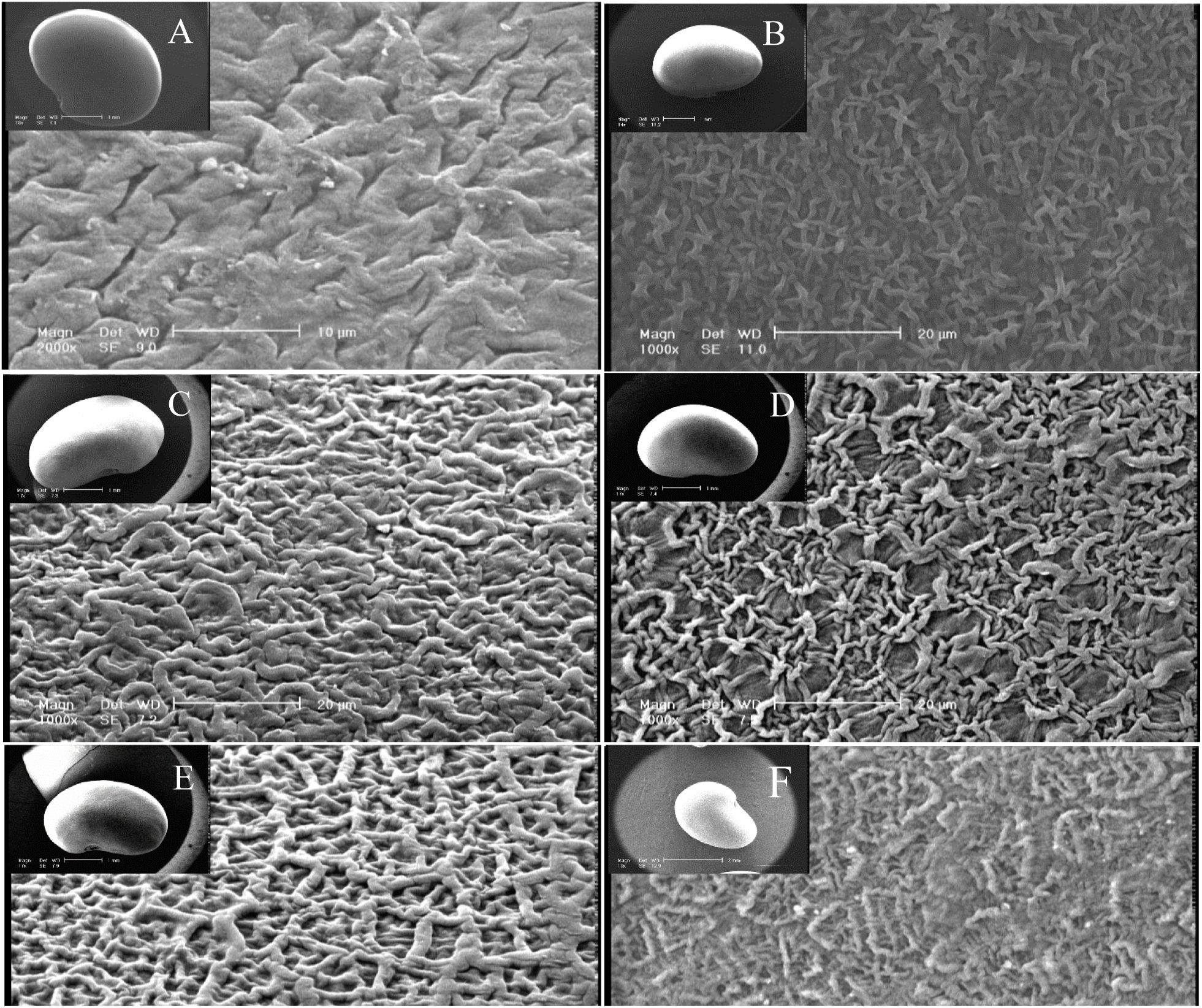
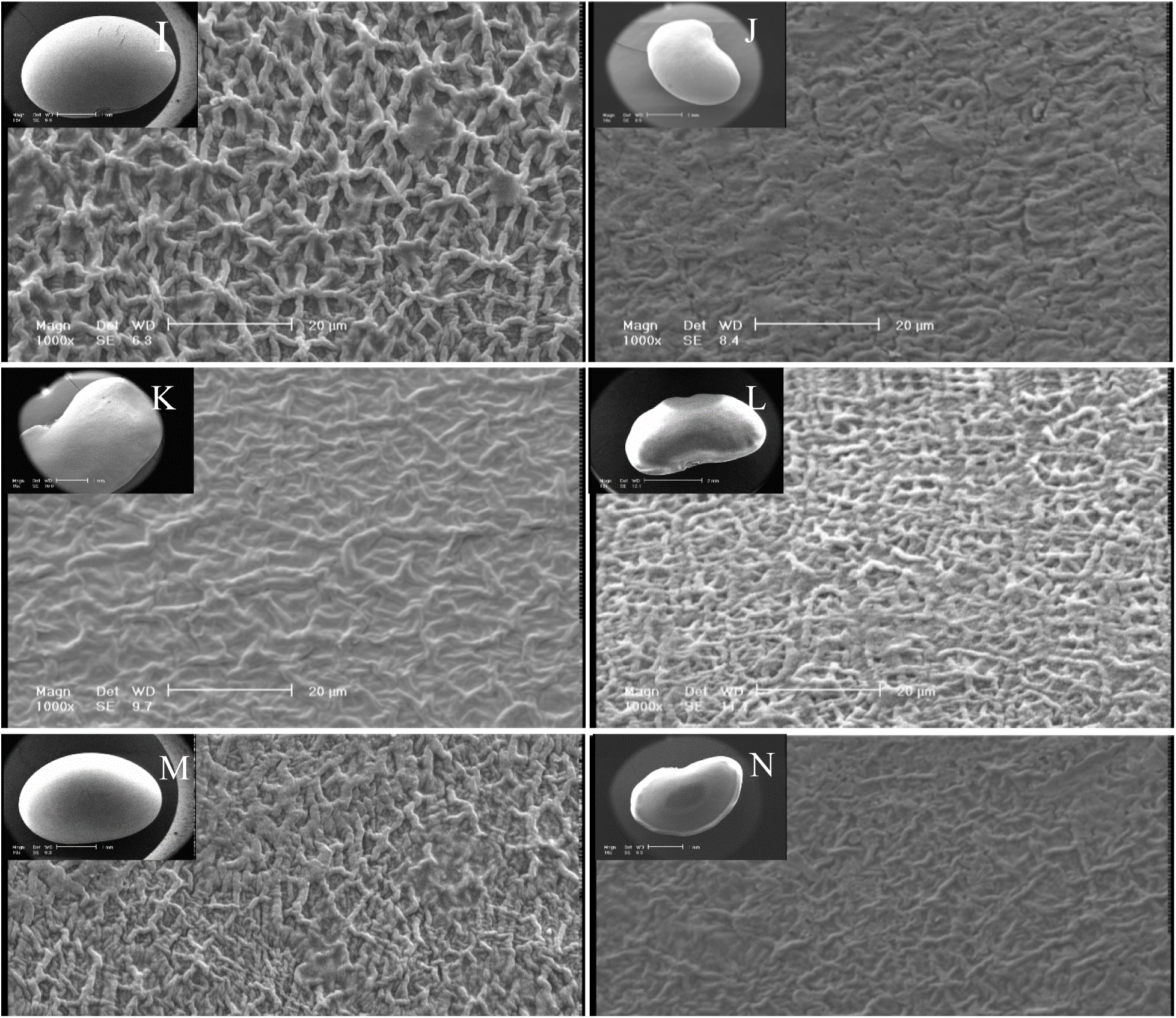
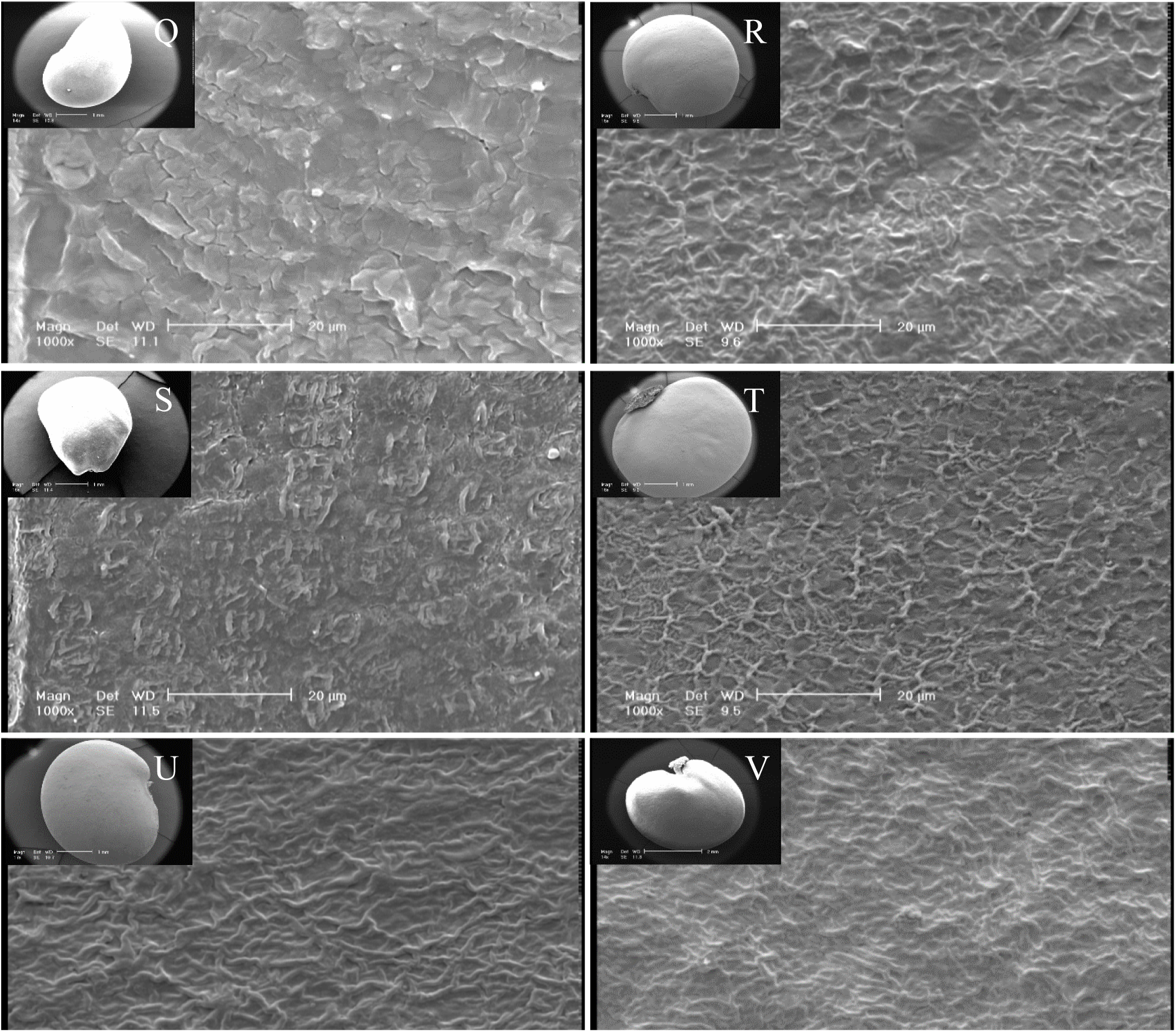
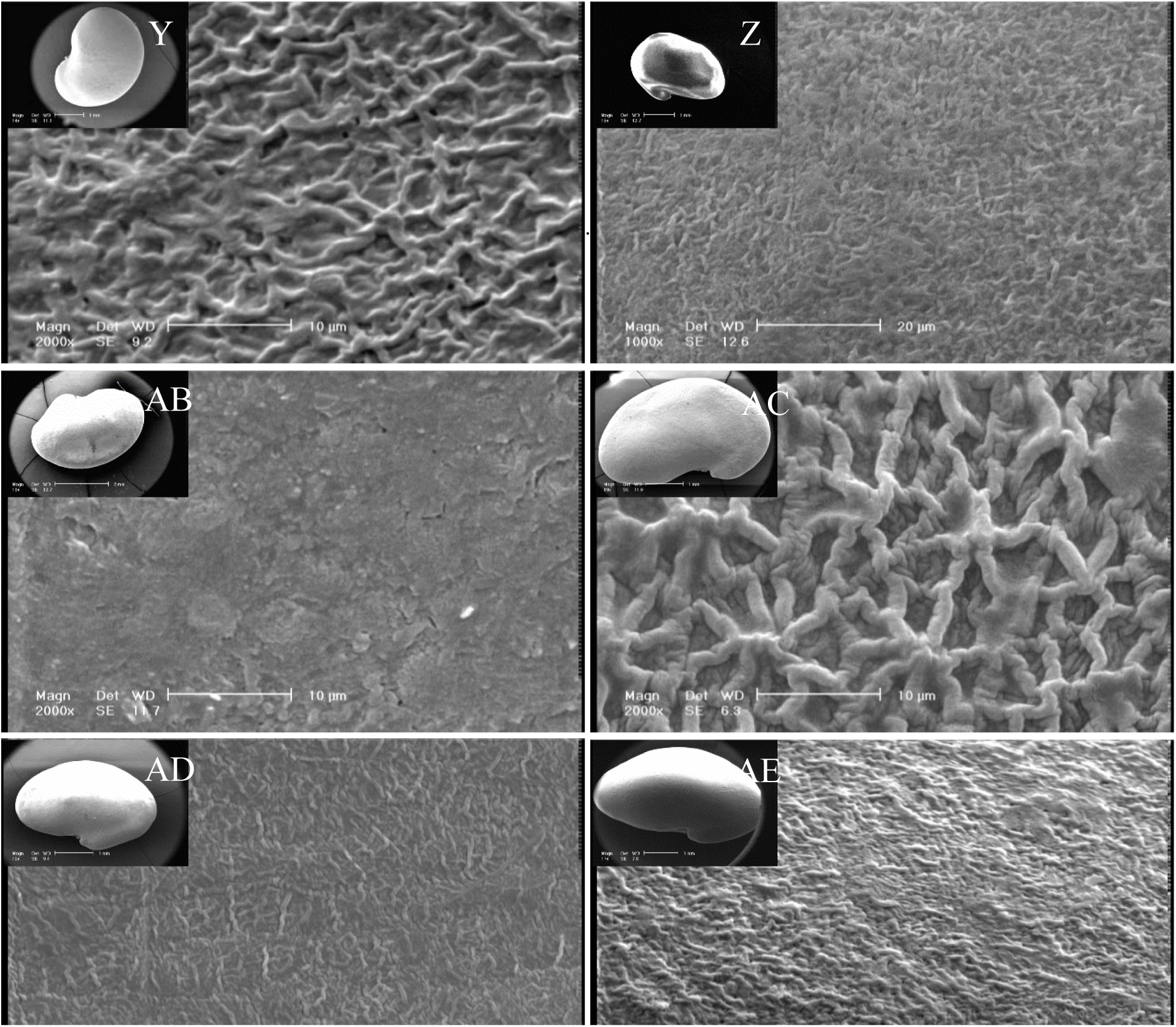
Scanning electron microscopy photographs of seed coat. A: *O. transcaucasia*, B: *O. arenaria*, C: *O. iberica*, D: *O. cyri-*Grossh, E: *O. altissima*, F: *O. alba*, G: *O. inermis*, and H: *O. petraea*. I: *O. oxyodonta*, J: *O. gracilis*, K: *O. persica*, L: *O. peduncularis*, M: *O. hajastana*, N: *O. megataphrose*, O: *O. montana*, and P: *O. viciifolia.* Q: *O. kemulariae*, R: *O. caput-galli*, S: *O. crista-galli*, T: *O. aequidentata*, U: *O. pulchella*, V: *O. melanotricha*, W: *O.argyrea*, and X: O. ptolemaica. Y: *O. hypargyrea*, Z: *O. michauxii*, AB: *O. sintenisii*, AC: *O. chorassanica*, AD: *O. bobrovii-Grossh*, AE: *O. pallasii*, and AF: *O. radiate*.

**Figure S5:**
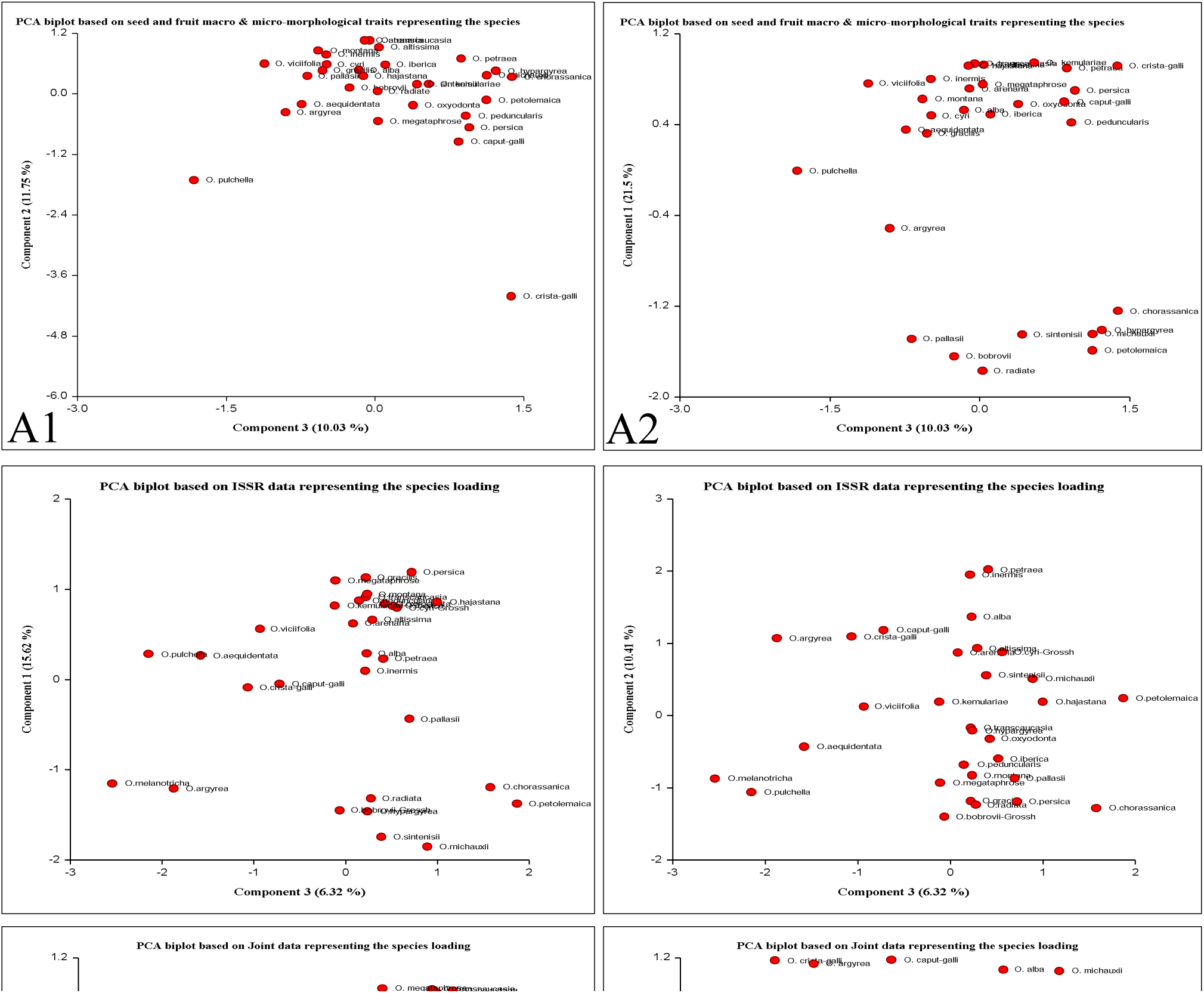
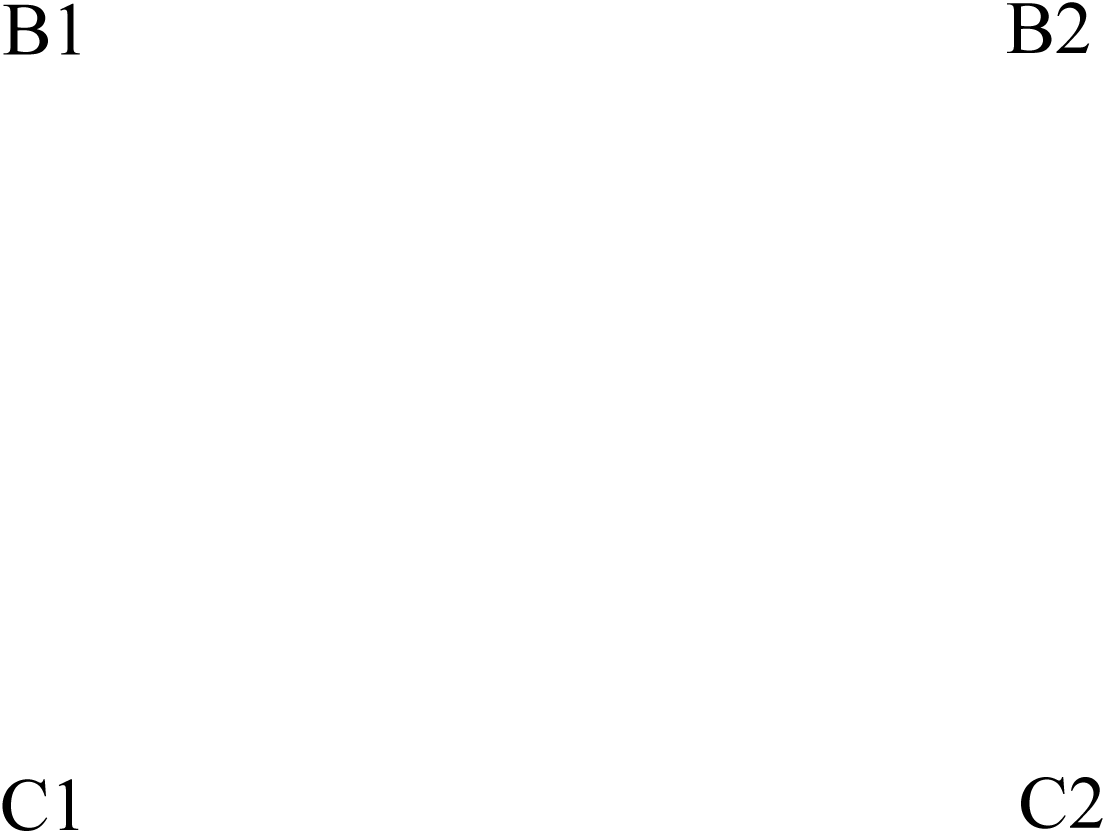
Two-dimensional representation of genetic relationships based on principal components analysis among the 31 *Onobrychis* species. A: seed and fruit macro & micro-morphological traits, B: PCA based on ISSR data, and C: based on joint data.

## References

1) Abou-El-Enain MM. 2002. Chromosomal criteria and their phylogenetic implications in the genus Onobrychis Mill, Sect. Lophobrychis (Leguminosae) with special reference to Egyptian species. Botanical Journal of the Linnean Society 139: 409–414.

2) Abou-El-Enain MM. 2004. SDS-PAGE of seed protein criteria in relation to taxonomy of *Onobrychis* sect. Lophobrychis s. str. and the Egyptian species. Cytologia 69: 351–358.

3) Al-Ghamdi FA, Al-Zahrani RM 2012. Seed morphology of some species of Tephrosia Pers. (Fabaceae) from Saudi Arabia identification of species and systematic significance. Feddes Repertorium 121: 59–65.

4) Amirahmadi A, Kazempour-Osaloo S, Kaveh A, Maassoumi AA, Naderi R. 2016. The phylogeny and new classification of the genus *Onobrychis* (*Fabaceae*-*Hedysareae*): evidence from molecular data. Plant Systematics and Evolution 302:1445–1456.

5) Arslan E, Ertugrul K. 2010. Genetic relationships of the genera Onobrychis, Hedysarum, and Sartoria using seed storage proteins. Turkish Journal of Biology 34: 67–73.

6) Attar F, Keshvari A, Ghahreman A, Zarre S, Aghabeigi F. 2007. Micromorphological studies on Verbascum (Scrophulariaceae) in Iran with emphasis on seed surface, capsule ornamentation and trichomes. Flora 202: 169–175.

7) Avcı A, Ilhan E, Erayman E, Sancak C 2014. Analysis of Onobrychis genetic diversity using SSR markers from related legume species. The Journal of Animal & Plant Sciences 24(2): 556–566

8) Avci S, Tekin N, Sancak C, Özcan S, Marang AO. 2016. Phylogenetic relationship of some Onobrychis taxa naturally grown in Turkey based on morphology and nuclear ribosomal DNA ITS Sequences. Legume Research 39(5): 665–67.

9) Ball PW 1968. Onobrychis Miller. In: Tutin TG, Heywood VH, Burges NA, Moore DM, Valentin SM (ed) Flora Europaea, Vol 2. Cambridge: Cambridge University, pp 187–191.

10) Bandara NL, Papini A, Mosti S, Brown T, Smith LMJ. 2013. A phylogenetic analysis of genus *Onobrychis* and its relationships within the tribe Hedysareae (Fabaceae). Turk J Bot 37: 981–992.

11) Barthlott W (1984). Microstructural features of seed surfaces. In: Heywood VH, Moore DM (Ed) Current concepts in plant taxonomy. London.

12) Bhattarai S, Coulman B, Fu YB, Beattie AD, Biligetu B 2018. Genetic diversity and relationship of sainfoin (*Onobrychis viciifolia* Scop.) germplasm as revealed by amplified fragment length polymorphism markers. Canadian Journal of Plant Science 98: 543–551

13) Cenci CA, Bassi G, Ferranti E, Romano B. 2000. Some morphometric, anatomical and biochemical characteristics of fruits and seeds of *Onobrychis* spp. in Italy. Plant Biosystems 134: 91–98.

14) Davis PH, Mill RR, Tan K. 1988. Flora of Turkey and the East Aegean Islands (Supplement), Vol 10. Edinburgh: Edinburgh University, pp 561–588.

15) Duan L, Wen J, Yang X, Liu PL, Arslan E, Ertuğrul K, Chang ZY. 2015. Phylogeny of *Hedysarum* and tribe Hedysareae (Leguminosae: Papilionoideae) inferred from sequence data of ITS, matK, trnL-F and psbA-trnH. Taxon 64 (1):49–64.

16) Earl DA. 2012. STRUCTURE HARVESTER: A website and program for visualizing STRUCTURE output and implementing the Evanno method. Conservation Genetics Resources 4(2), 359–361.

17) Elena T. 2006. Cytological aspects of the Onobrychis genus. Buletin USAMV 62: 154–158.

18) Emre I, Turgut-bahk D, Sahm A, Kursat M. 2007. Total Electrophoretic Band Patterns of Some Onobrychis Species Growing in Turkey. Journal of Agriculture and Environmental Sciences 2: 123–126.

19) Excoffier L, Lischer HEL. 2010. Arlequin suite ver 3.5: A new series of programs to perform population genetics analyses under Linux and Windows. Molecular Ecology Resources 10: 564–567.

20) Gontcharova SB, Gontcharov AA, Yakubov VV, Kondo K. 2009. Seed surface morphology in some representatives of the Genus Rhodiola sect. Rhodiola (Crassulaceae) in the Russian Far East. Flora 204: 17–24.

21) Hartl, D. L., Clark, A. G., & Clark, A. G. (1997). Principles of Population Genetics. Vol. 116. (Sinauer Associates,).

22) Hayot-Carbonero C. 2011. Sainfoin (Onobrychis viciifolia), a forage legume with great potential for sustainable agriculture, an insight on its morphological, agronomical, cytological and genetic characterization. PhD Thesis,, Univercity of Manchester, Manchester, U.K.

23) Hayot-Carbonero C, Carbonero F, Smith LMJ, Brown TA. 2012. Phylogenetic characterisation of Onobrychis species with special focus on the forage crop Onobrychis viciifolia Scop. Genetic Resources and Crop Evolution 59: 1777–1788.

24) Johnson LA, Huish KH, Porter JM. 2004. Seed surface sculpturing and its systematic significance in Gilia (Polemoniaceae) and segregated genera. International Journal of Plant Sciences 165: 153–172.

25) Karcz J, Ksiazazyk T, Maluszynska J. 2005. Seed coat patterns in rapid-cycling Brassica forms. Acta Biologica Cracoviensia s. Botanica 47: 159–165.

26) Khan MMH, Rafii MY, Ramlee SI, Josoh M, Mamum DA, Halidu J. 2021. DNA fingerprinting, fixation-index (Fst), and admixture mapping of selected Bambara groundnut (Vigna subterranea [L.] Verdc.) accessions using ISSR markers system. Scientific Reports 11:14527. https://doi.org/10.1038/s41598-021-93867-5

27) Koul KK, Nagpal R, Raina SN. 2000. Seed Coat Microsculpturing in Brassica and Allied Genera (Subtribes Brassicinae, Raphaninae, Moricandiinae). Annals of Botany 86: 385–397.

28) Moazzeni H, Zarre S, Al-Shehbaz IA, Mummenhoff K. 2007. Seed-coat microsculpturing and its systematic application in Isatis (Brassicaceae) and allied genera in Iran. Flora 202: 447–454.

29) Murray MG, Thompson WF. 1980. Rapid isolation of high molecular weight plant DNA. Nucleic Acids Research 8: 4321–4325.

30) Pavlova DK, Manova VI. 2000. Pollen morphology of the genera Onobrychis and Hedysarum (Hedysarea, Fabaceae) in Bulgaria. Annales Botanici Fennici 37: 207–217.

31) Powell W, Morgante M, Ander C, Hanafey M, Vogel J, Tingy S, Rafalaski A. 1996. The comparision of RFLP, RAPD, AFLP and SSR (microsatellite) marker for germplasm analysis. Molecular Breeding 2: 225–238.

32) Pradeep Reddy M, Sarla N, Siddiq E. 2002. Inter simple sequence repeat (ISSR) polymorphism and its application in plant breeding. Euphytica 128: 9–17. https://doi.org/10.1023/A:1020691618797

33) Prevost A, Wilkinson MJ. 1999. A new system of comparing PCR primers applied to ISSR fingerprinting of potato cultivars. Theoretical and Applied Genetics 98: 107–112.

34) Pritchard JK, Wen W, Falush D. 2010. Documentation for STRUCTURE Software: Version 2 (University of Chicago).

35) Punt W, Blackmore S, Nilsson S, Thomas A. 1994. Glossary of pollen and spore terminology. LPP Foundation, Utrecht.

36) Rechinger KH. 1969. Flora Iranica. In: Reshinger KH (ed). Vol. 157. Akademische Druck-und Verlgsantalt, Graz, Austria. p. 387.

37) Rohlf FJ (1998). NTSYS-pc numerical taxonomy and multivariate analysis system. Version 2.00. Exeter Software, Setauket, NY.

38) Roldan-Ruiz I, Dendauw J, Bockstaele EV, Depicker A, Loose MD. 2000. AFLP markers reveal high polymorphic rates in ryegrasses (*Lolium* spp.). Molecular Breeding 6: 125–134.

39) Safaei Chaei Kar S, Ghanavati F, Naghavi MR, Amirabadi-zade H, Rabiee R. 2014. Molecular phylogenetics of the Onobrychis genus (Fabaceae: Papilionoideae) using ITS and trnL–trnF DNA sequence data. Australian Journal of Botany 62: 235–250.

40) Stearn WT (1983). Botanical Latian. Nelson. London.

41) Tamura K, Stecher G, Peterson D, Filipski A, Kumar S. 2013. MEGA6: Molecular evolutionary genetics analysis version 6.0. Molecular Biology and Evolution 30(12): 2725–2729.

42) Toluei Z, Atri M, Ranjbar M, Wink M. 2013. Iranian Onobrychis carduchorum (Fabaceae) populations: morphology, ecology and phylogeography. Plant Ecology and Evolution 146: 53– 67

43) Vural C, Ekici M, Akan H, Aytac Z. 2008. Seed morphology and its systematic implications for genus Astragalus L. sections Onobrychoidei DC., Uliginosi Gray and Ornithopodium Bunge (Fabaceae). Plant Systematics and Evolution 274: 255–263 .

44) Yeh FC, Yang R, Boyle T. 1999. Popgene version 1.32: Microsoft Windows-based freeware for population genetic analysis. University of Alberta, Edmonton.

45) Yildiz B, Ciplak B, Aktoklu E. 1999. Fruit morphology of sections of the genus Onobrychis Miller (Fabaceae) and its phylogenetic implications. Israel Journal of Plant Sciences 47: 269–282.

46) Zarei A, Erfani-Moghadam J. 2021. SCoT markers provide insight into the genetic diversity, population structure and phylogenetic relationships among three Pistacia species of Iran. Genetic Resources and Crop Evolution. 68(4), 1625–1643.

47) Zarrabian M, Majidi M, Ehtemam MH. 2013. Genetic Diversity in a Worldwide Collection of Sainfoin Using Morphological, Anatomical, and Molecular Markers. Crop Science 53: 1–14.

48) Zarrabian, M, Majidi, M.M. (2015). Genetic diversity and relationships within and among *Onobrychis* species using molecular markers. Turkish Journal of Botany. 39: 681–692.

